# Predicting the protein interaction landscape of a mycobacterial pathogen

**DOI:** 10.64898/2026.07.15.738315

**Authors:** Horia Todor, Lili M. Kim, Evan Billings, Anna E. Grzegorzewicz, Hannah N. Burkhart, Michael A. DeJesus, Allen Na, Samuel Nitz, Scarlet S. Shell, Elizabeth A. Campbell, Mary Jackson, Filippo Mancia, Jeremy M. Rock, Carol A. Gross, James Chen

## Abstract

High-dimensional phenotypic screens of bacterial loss-of-function mutant libraries have determined gene-gene connections and specific phenotypes for thousands of bacterial genes in many species, but deciphering the underlying mechanisms remains decidedly low-throughput. Here, we demonstrate the utility of proteome-wide AI-based protein-protein interaction (PPI) predictions for overcoming this gap by using pooled-AlphaFold3 to assess all ∼1.3 million possible pairwise interactions in the proteome of *Mycobacterium leprae.* We identify ∼2,000 strong and intermediate PPIs that underlie a significant fraction of phenotypes and gene-gene connections observed in large-scale chemical genomics screens from *Mycobacterium tuberculosis, Mycobacterium smegmatis,* and *Corynebacterium glutamicum*. This combined approach predicts specific functions for dozens of previously uncharacterized core, conserved, and essential mycobacterial proteins. We highlight new information derived from the study, including insights into mycobacterial envelope assembly, peptidoglycan remodeling, and new modulators of the central dogma enzymes RNA polymerase and DNA gyrase. These data establish combined pooled-AlphaFold3 PPI prediction and high-throughput genomics approach as the gold standard for large-scale characterization of protein function.

## INTRODUCTION

Bacteria are central drivers of terrestrial, aquatic, and host-associated ecosystems, and have major impacts on human health and disease. Despite this, our understanding of bacteria is strikingly incomplete. Since the first bacterial genome was sequenced more than 30 years ago, we have been accumulating genome sequences encoding genes of completely or partially unknown function. That ∼⅓ of all bacterial genes are uncharacterized greatly limits our ability to understand and manipulate bacteria^1^. High-throughput functional genomics screens using genome-wide knockout or knockdown libraries have partially closed this gap, allowing the determination of gene-inactivation phenotypes at scale in diverse bacteria^2–4^. However, deciphering the function of a gene from its phenotypes remains difficult, especially for truly novel genes such as those with domains of unknown function (DUFs), limiting the impact of these high-throughput studies.

Protein-protein interactions (PPIs) are at the center of almost all major biological processes and are thus the ultimate currency for determining gene function. We recently developed pooled-AlphaFold3^5^, a strategy that reduces the high false-positive rate of AI-structure based approaches by co-folding ∼10-15 unrelated proteins, thereby enabling accurate proteome-wide (all-by-all) PPI predictions. Here we show how combining predicted PPIs with experimentally derived phenotypic correlations can functionally and structurally associate uncharacterized genes with known partners, thus uncovering gene functions and facilitating the discovery of their precise mechanistic role.

We demonstrate this combined approach by applying it to Mycobacteria, a diverse bacterial clade characterized by an atypical and impermeable cell envelope. Despite the central role mycobacteria play in human health and disease (e.g., *Mycobacterium tuberculosis*, the world’s leading infectious disease killer), they encode dozens of conserved essential genes of unknown function, many of which are involved in the synthesis of their atypical cell envelope^6–8^. We use pooled-AlphaFold3 to predict all pairwise PPIs in the proteome of *Mycobacterium leprae*^9^ and combine these PPI predictions with high-throughput phenotypic data from related organisms^10–12^. We chose to predict PPIs in *M. leprae* because its minimal proteome (1,610 proteins) with ∼1.3M unique pairwise interactions: 1) reduces the computational requirements compared to *M. tuberculosis* (∼7M unique pairs) or *Mycobacterium smegmatis* (∼22M unique pairs); 2) reduces the number of false positive interactions (which scale with the number of protein pairs) especially for homologous genes that may have partially or completely overlapping multiple sequence alignments (MSAs)^13^; 3) retains almost all essential and conserved mycobacterial processes^14^.

We uncover functions for numerous essential genes and novel partners for essential processes. First, we identified new players and interactions in the regulation, synthesis, and synchronization of mycomembrane components^8^. We predicted and verified a role for EphE in the synthesis of phosphatidyl-myo-inositol mannosides (PIMs, major constituents of the mycobacterial inner membrane^15^), associated two essential proteins with MptB (which is involved in the mannosylation of PIMs^16^), and identified a probable regulatory connection between arabinogalactan (AG) synthesis and mycolic acid synthesis. Second, we found functional partners for core mycobacterial enzymes that revealed unexpected similarity to eukaryotic complexes. For example, we identified a heteromeric protein mannosyltransferase complex that functionally resembles eukaryotic complexes and uncovered a Trm112p-family methyltransferase activator involved in the regulation of mycolic acid export. Finally, we identify new essential and non-essential partners of RNA-polymerase and DNA-gyrase, core central dogma enzymes and the targets of first- and second-line antibiotics in *M. tuberculosis*^17^. This data will nucleate future functional studies in mycobacteria and the structures of putative interactions can be mined to generate both specific and general hypotheses about how proteins interact. Together, our data establishes a combined proteome-wide PPI prediction and high-throughput chemical genomics approach as the gold standard for functional characterization of bacterial protein function at scale.

## RESULTS

### A proteome-wide PPI map in *M. leprae*

We used pooled-AlphaFold3^5^ to predict the protein-protein interaction network of *M. leprae* (Figure 1A). The *M. leprae* genome encodes 1,610 proteins. To query all ∼1.3M pairwise interactions, we designed a set of 17,861 pools such that each unique pair of proteins appeared in at least one pool (Table S1). The pools consisted of 2 to 27 proteins (median n = 14) whose sizes summed to ∼5,000 amino acids (aa) (Figure S1A-B). All 17,861 pools were successfully run on the free AlphaFold3 server (596 person-days of free daily quota). For each pool we averaged the matrix of pairwise interface predicted template modeling (ipTM) scores for all 5 diffusion samples and corrected for the size-bias in ipTM scores as previously described^5^. A total of 378,245 protein pairs appeared in more than one pool and we used these pairs to assess reproducibility of ipTM scores (r = 0.59; Figure S1C). Reproducibility was consistent with our previous proteome-wide pooled-AlphaFold3 study^5^. We used pairwise scores to construct a 1,609 x 1,609 matrix of pairwise size-corrected ipTM scores (Table S2). As expected considering the known scarcity of PPIs^18^, 98.2% of interactions exhibited a size-corrected ipTM < 0.05. Our study identified 406 strong (size-corrected ipTM > 0.4) interactions, 1,759 intermediate (size-corrected ipTM > 0.2 & < 0.4) interactions, and 21,074 marginal (size-corrected ipTM > 0.05 & < 0.2) interactions (Figure 1B). These interactions involved almost every protein: 28% of the proteome exhibited at least 1 strong interaction, 68% had at least 1 intermediate interaction, and 98% had at least 1 weak interaction (Figure 1C).

**Figure 1.**
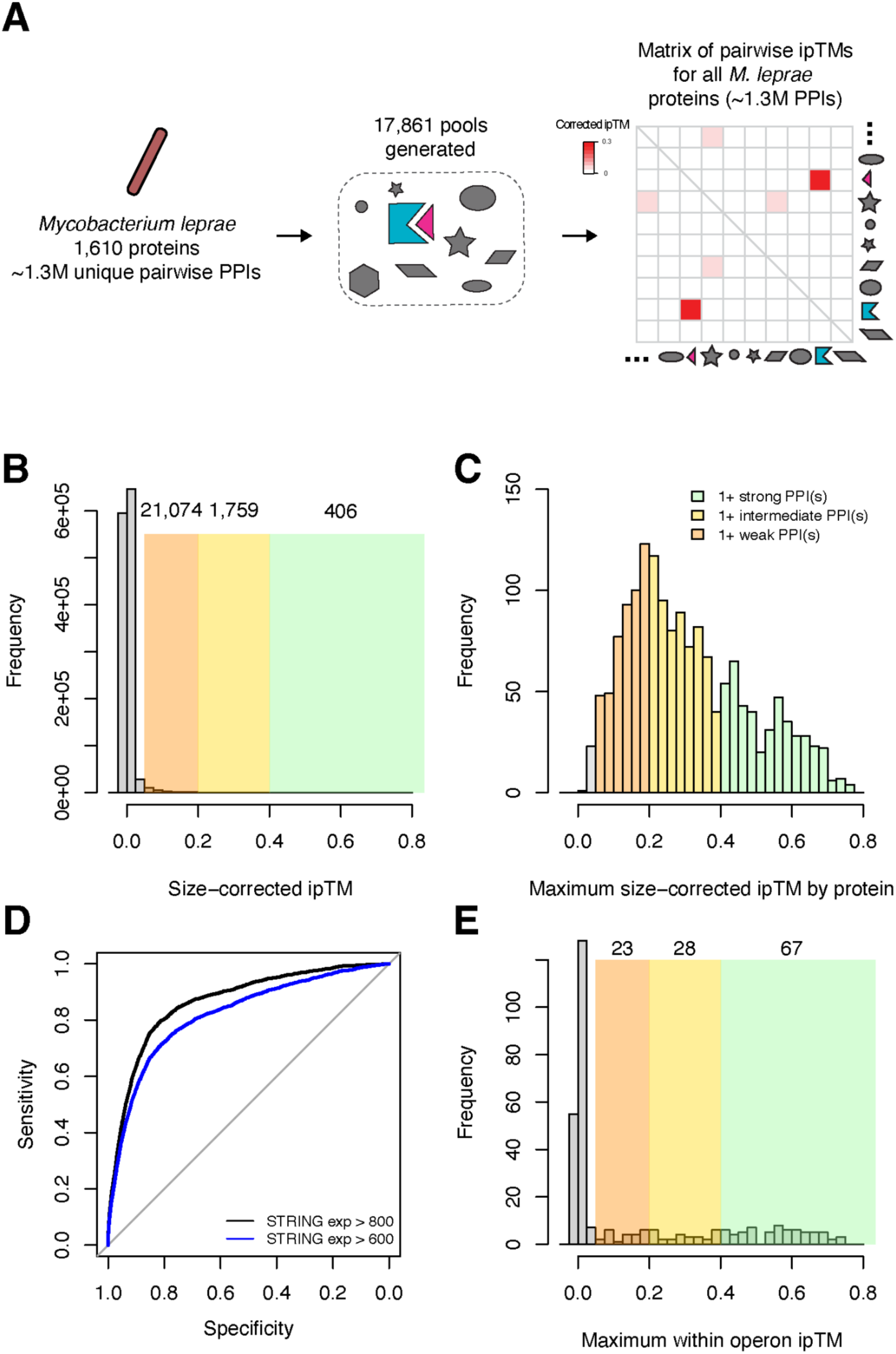
**(A)** Schematic of Pooled-AlphaFold3 approach for genome-wide prediction of PPI in *M. leprae*. **(B)** Distribution of size-corrected pairwise ipTM scores for all 1,293,636 queried protein pairs from the *M. leprae* genome. The background of the histogram is colored green, yellow, and red to indicate the region containing strong, intermediate, and weak interactions (respectively). **(C)** Most proteins were predicted to make at least one weak, intermediate, or strong interaction. Histogram shows the strongest predicted interaction for each of the 1,609 queried proteins. Colors are as in (B). **(D)** ROC curves showing that predicted PPI accurately capture known interaction in the STRING experimental database. ROC curve plots sensitivity (true positive rate) on the y-axis against specificity (1-false positive rate) on the x-axis across all thresholds of the predictor (in this case, size-corrected ipTM). AUROC for STRING experimental > 800 = 0.86; black. AUROC for STRING experimental > 600 = 0.82; blue. **(E)** Histogram of maximum size-corrected ipTM within the 309 operons in *M. leprae*. Background colors are as in (B).

We used two metrics to assess the accuracy and utility of our predictions. First, we benchmarked our predicted interactions against the STRING database “experimental” channel, which consists of protein–protein association evidence imported from repositories such as BioGRID, PDB, and IntAct rigorously propagated through bacterial phylogeny to appropriately integrate information from related species^19^. Only 2,877 protein pairs (0.2%) had strong evidence of interaction (score >800) in this database. We found that our pooled-AlphaFold3 data accurately predicted these known interactions, as quantified by an area under the receiver operating characteristic curves (AUROC) of 0.86 for STRING-experimental scores >800 (Figure 1D). Second, we assessed the frequency of predicted PPIs between proteins encoded in the same operon. Since operons are frequently mono-functional, we expect proteins encoded within the same operon to frequently interact. However, since this genomic information is not passed to AlphaFold3 (which sees only protein sequences), the frequency of predicted PPIs between genes encoded in the same operon is a relatively unbiased metric of overall PPI prediction performance. We found that of the 309 operons encoding at least two proteins (i.e., not pseudogenes) in *M. leprae*^20^, 67 (21.7%) had at least one strong predicted interaction between their encoded proteins, and protein pairs encoded by the same operon were >300-fold more likely to have a strong predicted interaction than a pair of proteins chosen at random (Figure 1E). Together, these metrics suggest that our pooled-AlphaFold3 dataset of predicted *M. leprae* PPIs accurately identifies interacting protein pairs.

### *M. leprae* predicted PPIs recapitulate known but structurally uncharacterized PPIs

While many of our strong hits were either located in the same operon (23%), or had strong evidence of interaction in the STRING-experimental channel (21%), a majority of strong predicted interactions were potentially novel (e.g., STRING-experimental score = 0). To ascertain whether these predicted interactions represent true PPIs, we searched for evidence of physical interaction for these protein pairs in the mycobacterial literature. We found many of our “new” interactions had been recently described in other mycobacteria using either targeted pull-downs or in structures published after the AlphaFold3 training cutoff date. For example, we predicted the recently described tripartite interaction between the essential arabinosyltransferase AftB and the β-propeller ML1720-homolog (size corrected ipTM = 0.52; Table S2), and between ML1720 and the periplasmic binding protein FecB (size corrected ipTM = 0.49; Table S2) as well as the interaction between AftA and LpqZ (size corrected ipTM = 0.31; Table S2) and between AftC and AftH^10,21,22^ (size corrected ipTM = 0.21; Table S2). Similarly, we predicted the recently described interaction between Mce1C and ML0520c/LucB^23^ (size corrected ipTM= 0.47; Table S2), the structurally uncharacterized interaction between MurA and its regulator CwlM^24^ (size corrected ipTM = 0.58; Table S2), the interaction between HtrA and LppZ^25^ (size corrected ipTM = 0.37; Table S2), and numerous other validated interactions that are not yet represented in STRING (Table S2).

In addition to capturing many recently reported physical interactions, our predicted PPIs also suggested that many previously reported functional interactions are underpinned by direct physical interactions. For example, LmeA (ML2195) has previously been reported to be important for the activity and stability of MptA (ML0899)^26^, the major mannan polymerase in mycobacteria. Our data predicts that this is due to a direct interaction between MptA and LmeA (size corrected ipTM = 0.61; Table S2). Similarly, LpqW (ML1497c) has been functionally linked to MptB (ML0591c)^27^ and our data suggests a direct interaction (size corrected ipTM = 0.50; Table S2). Finally, ML1816, a member of the antibiotic resistance ABC-F protein family, rescues linezolid bound ribosomes^12,28^. Our data predicts a direct interaction with RplA (size corrected ipTM = 0.32; Table S2), one of the major rRNA binding proteins, illuminating the mechanism of ML1816’s interaction with the ribosome. Together, these data suggest that our pooled-AlphaFold3 data can identify functionally important PPIs.

### PPIs often underlie phenotypic correlations in high-throughput chemical genomics screens

The fact that many functional interactions between proteins in *M. smegmatis, M. tuberculosis,* and *C. glutamicum* proteins could be explained by PPIs between their *M. leprae* homologs suggested that many of the *M. leprae* PPIs are conserved, and may explain other previously observed functional interactions in related organisms. Mycobacteria and related bacteria have been subject to numerous high-throughput genome-wide phenotypic screens^29^, most recently using CRISPRi approaches^10,12,30^, which allow assessment of essential gene depletion phenotypes (in addition to non-essential gene depletion phenotypes). Such screens can functionally associate proteins based on shared phenotypes across multiple conditions (phenotypic signatures)^10,31^, raising the question of the extent to which phenotypic correlations between genes can be explained by the predicted *M. leprae* PPIs.

To answer this question, we first mapped *M. leprae* genes onto their homologs in *M. tuberculosis* (homologs for 1429/1610 *M. leprae* proteins), *M. smegmatis* (homologs for 1322/1610 *M. leprae* proteins), and *C. glutamicum* (homologs for 951/1610 *M. leprae* proteins) (Table S3): three organisms which have been extensively studied using broad chemical genomics screens^10–12,29^. We then compared the phenotypic correlations between genes with the size-corrected ipTM of their *M. leprae* homologs to identify physically interacting co-functioning proteins (Figure 2A-C). Phenotypically correlated gene pairs were heavily enriched in putatively interacting protein pairs (Table S4). For example, the most highly correlated gene pairs (r > 0.8) were highly enriched for protein pairs with size-corrected ipTM > 0.4 (50-fold in *M. tuberculosis* (Figure 2A), 367-fold in *M. smegmatis* (Figure 2B), and 226-fold in *C. glutamicum* (Figure 2C). The differences in fold enrichment can be attributed to evolutionary distance between each organism and *M. leprae*, different methodologies (e.g. Tn-seq vs CRISPRi), and the number and diversity of chemicals tested (9 drugs in *M. tuberculosis* compared to ∼30 for *M. smegmatis* and *C. glutamicum*). Intermediate and strong predicted PPIs explained a substantial fraction of highly phenotypically correlated genes in *M. smegmatis* (18%) and *C. glutamicum* (12%), suggesting that PPIs frequently underlie phenotypic correlations (Table S4) and that these PPIs can be accurately assessed using our method.

**Figure 2.**
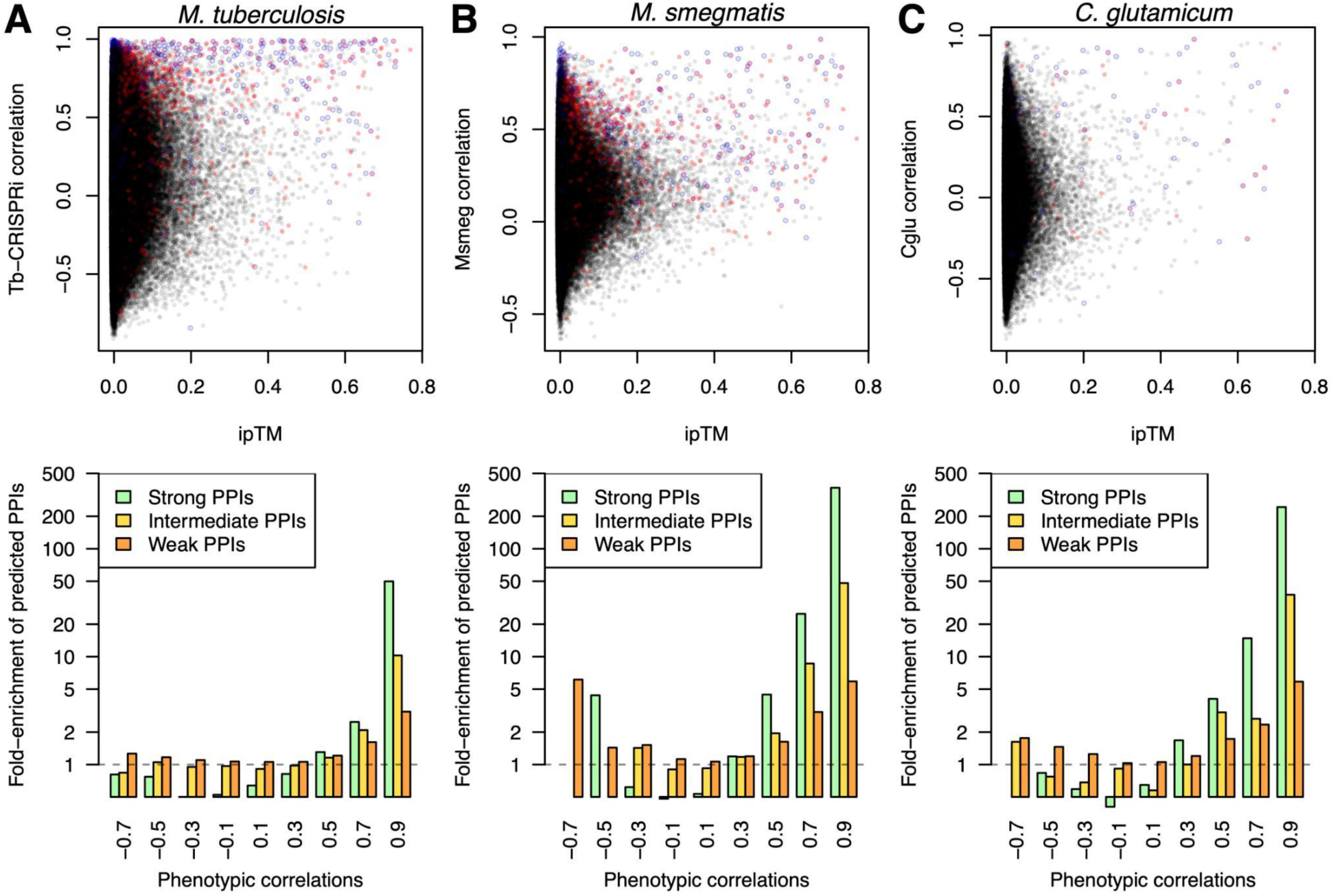
Predicted PPIs explain phenotypic correlations in diverse chemical genomics data sets. Size-corrected ipTM of all protein pairs in *M. leprae* plotted against the correlation between the knockdown phenotypes of their homologs (if present) in the *M. tuberculosis* CRISPRi chemical genomics dataset (A), the *M. smegmatis* CRISPRi chemical genomics dataset (B), or the *C. glutamicum* TnSeq chemical genomics dataset (C). Interactions with strong evidence of interaction in STRING (experimental channel > 800) are highlighted in red. Gene pairs in the same operon are bordered in blue. Histograms below show the relative enrichment of strong (green), intermediate (yellow), and weak (red) predicted PPIs among gene pairs with correlations in 0.2 width bins centered on the labels (i.e., *M. smegmatis* gene pairs with correlations between 0.8 and 1 are 367-fold enriched in strong predicted PPIs relative to background).

### A global network of functionally important predicted PPIs

The new connections revealed by our data involve almost every aspect of mycobacterial biology, including respiration, protein secretion, metabolism, and translation, and especially the mycomembrane. This is likely due both to the mycobacterial-specific nature of these genes, and to the chemicals chosen for the *M. smegmatis* and *C. glutamicum* chemical genomics screens, both of which focused on antibiotic and chemical stressors affecting the mycomembrane. Below, we demonstrate the insights that can be revealed by this approach by analyzing and validating 8 novel complexes involved in diverse aspects of mycobacterial biology. We focused predominantly on essential, conserved, or otherwise important complexes encoded at distinct genomic loci (i.e., not in an operon), since these are the most impactful and unexpected. Our first five vignettes focus on various parts of the mycobacterial cell envelope, uncovering new players in early and late PIM synthesis, the coordination and regulation of mycolic acid export, and peptidoglycan remodeling. We then describe the discovery and validation of a heteropentameric protein complex that mannosylates periplasmic proteins, and end by uncovering small (but important) proteins that likely modulate the activity of RNA polymerase and DNA gyrase.

### EphE (ML2297) and ML0451c function with PimA in PIM synthesis

PIMs are essential glycolipids in mycobacteria and other actinobacteria. The PIM synthesis pathway (depicted in Figure 3A) begins with the synthesis of phosphatidyl-inositol phosphate (PIP) by PgsA^32,33^. PIP is subsequently dephosphorylated by an as-yet unidentified phosphatase to generate phosphatidylinositol (PI)^33,34^, which is then mannosylated by the GT-4 glycosyltransferase PimA to produce PIM_1_^35^. PIM_1_ is acetylated and further mannosylated to generate Ac_1_PIM_2_ and Ac_2_PIM_2_, the principal glycolipid constituents of the inner leaflet of the mycobacterial inner membrane^15^. A portion of Ac_2_PIM_2_ is subsequently translocated to the outer leaflet, where it serves as the precursor for higher-order PIMs, lipomannan (LM), and lipoarabinomannan (LAM).

**Figure 3.**
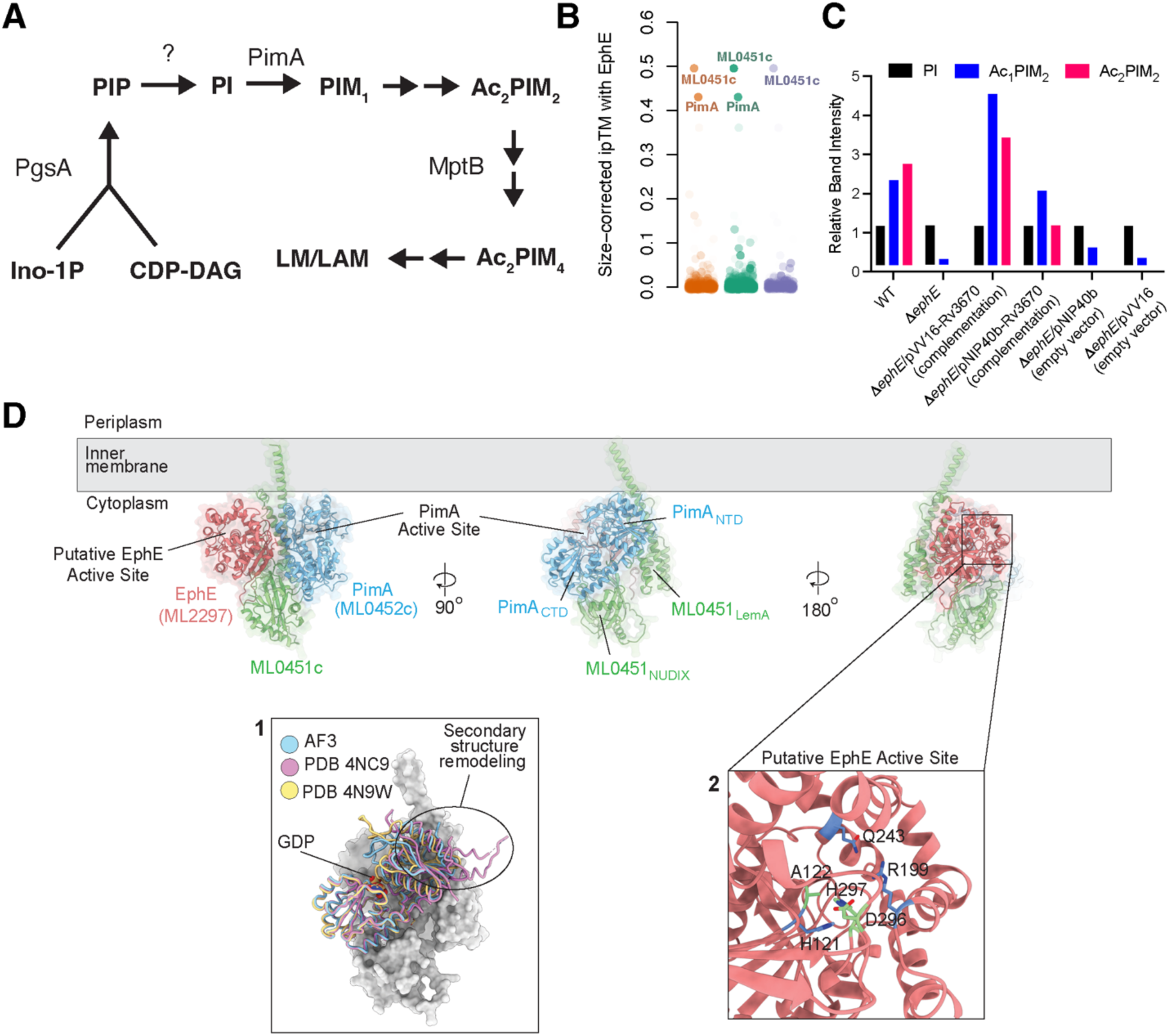
EphE is a novel enzyme involved in PIM synthesis in mycobacteria. **(A)** Schematic of PIM synthesis in Mycobacteria. **(B)** Size-corrected ipTMs between EphE (ML2297/MSMEG6184/Rv3670/Cg0358) and all other proteins in the *M. leprae* data. Opacity of points represents the phenotypic correlations in *M. smegmatis* (orange), *M. tuberculosis* (green), and *C. glutamicum* (purple). ML0451c (MSMEG2963/Rv2609c/Cg1875) and ML0452c (PimA: MSMEG2935/Rv2610c/Cg1876) are the only EphE interactors with size-corrected ipTM > 0.4, and are highly phenotypically correlated in all datasets. **(C)** Relative radioactivity incorporated into PI, Ac₁PIM₂, and Ac_2_PIM₂ in total [1,2-^14^C]acetate-derived lipids prepared from *M. smegmatis* WT, *ΔephE*, *ΔephE* mutants complemented with *M. tuberculosis ephE* (*rv3670*) expressed from a replicative plasmid under the control of the *hsp60* promoter (pVV16-Rv3670) or from an integrative plasmid under control of its native promoter (pNIP40b-Rv3670) (“complementation”), and empty vector controls (*ΔephE/*pVV16 and *ΔephE*/pNIP40b, “empty vector”). Strains were analyzed by TLC using CHCl_3_/CH_3_OH/H_2_O (65:25:4, by vol) as the eluent (Figure S2E). Radioactivity was semi-quantified using a Sapphire Biomolecular Imager and the results are expressed relative to the radioactivity incorporated into PI by each strain arbitrarily set to 1 (Methods). PI, phosphatidy-*myo*-inositol; Ac₁PIM₂, triacylated phosphatidylinositol dimannosides; Ac₂PIM₂, tetraacylated phosphatidylinositol dimannosides. **(D)** AlphaFold3 model of the *Mycobacterium leprae* PimA–ML0451c–EphE complex. PimA (ML0452c, blue), ML0451c (green), and EphE (ML2297, salmon) are shown as ribbon representations overlaid with transparent molecular surfaces and positioned relative to the inner membrane (grey). The PimA catalytic pocket and a putative EphE active site are indicated. Orthogonal views obtained by 90° and 180° rotations reveal the spatial organization of the complex, with the N-terminal NUDIX domain and C-terminal LemA-like domain of ML0451c bridging PimA and EphE at the cytoplasmic face of the membrane. **(Inset 1)** Structural comparison of PimA in the AlphaFold3-predicted PimA–ML0451c–EphE complex (cyan) with the crystal structures of apo PimA (PDB: 4NC9^37^, magenta) and GDP-bound PimA (PDB: 4N9W^37^, yellow). Guanosine diphosphate (GDP) is shown as sticks. Within the predicted complex, PimA adopts a conformation resembling the GDP-bound state. The secondary-structure rearrangement observed in apo PimA would sterically clash with ML0451c, suggesting that complex formation stabilizes the closed, GDP-bound-like conformation. **(Inset 2)** Close-up view of the putative EphE active site predicted by AlphaFold3. Conserved catalytic residues H121, A122 and R199 (green), together with putative phosphate-binding residues Q243, D296 and H297 (blue), are shown as sticks and define a solvent-accessible cavity within the cytoplasmic domain of EphE. Notably, EphE lacks the conserved nucleophilic residue (Asp or Ser) typically found at the highly conserved nucleophile elbow, with alanine occupying position 122 instead.

Our pairwise predicted PPI data suggested that PimA (ML0452c) forms a complex with EphE (ML2297) and a membrane protein of unknown function, ML0451c (Figure 3B, Table S2). The association between EphE and early PIM synthesis is supported by genetic evidence showing that *ephE* mutants, but not mutants of other annotated epoxide hydrolases, phenocopy *pimA* mutants and mutants in other genes of the *pgsA*/*patA*/*pimA* operon in *M. tuberculosis, M. smegmatis, and C. glutamicum* (Figure S2A–C). To investigate whether EphE plays a role in PIM biosynthesis, we generated a *ΔephE* mutant in *M. smegmati*s (Figure S2D). The mutant exhibited impaired growth but remained viable, enabling analysis of membrane lipid composition by thin-layer chromatography (TLC). Strikingly, *ΔephE* cells displayed dramatically reduced levels of Ac_1_PIM_2_ and Ac_2_PIM_2_, the major PIM species in mycobacteria, indicating a critical role for EphE in PIM production (Figure 3C, Figure S2E).

To better understand what role EphE and ML0451c may play in early PIM synthesis, we co-folded the heterotrimeric complex. In the predicted structure, ML0451c occupies a central position between PimA and EphE at the cytoplasmic face of the inner membrane, with the active sites of both EphE and PimA facing towards the membrane (Figure 3D, left panel).

ML0451c, which is encoded at the distal end of the conserved *pgsA*/*patA*/*pimA* operon, contains a LemA-like domain (conserved in all PIM producing actinobacteria) and a C-terminal NUDIX hydrolase domain (specific to Mycobacteriaceae and Nocardiaceae). Its LemA and NUDIX domains form extensive interactions with PimA (Figure 3D, middle panel) and EphE (Figure 3D, right panel), suggesting that ML0451c functions as a membrane-associated scaffold, potentially coordinating substrate presentation and catalysis. LemA-like domains typically form four-helix bundles in which the first helix is substantially longer than the remaining helices and have been implicated in membrane remodeling processes in other bacteria (e.g., *Magnetospirillum magneticum*^36^). Consistent with this role, the AlphaFold3 model positions the extended first helix of ML0451c adjacent to the membrane, where it may potentially perturb local phospholipid packing and facilitate access of the active sites of PimA and EphE to phosphatidylinositol-containing substrates (Figure 3D, middle panel).

Previous structural studies demonstrated that PimA catalysis requires substantial conformational rearrangements involving both its flexible interdomain region and the relative orientation of its N-terminal and C-terminal Rossman-fold domains^37^. To determine if these conformational rearrangements are affected by interactions with EphE and ML0451c, we compared the structure of PimA within the predicted complex with the available crystal structures of apo PimA and GDP-bound PimA^37^. We found that in the putative complex, PimA adopts a conformation closely resembling the GDP-bound state (Figure 3D, inset 1). Remarkably, the secondary-structure rearrangement observed in apo PimA would sterically clash with ML0451c in the predicted complex, suggesting that complex formation may stabilize a closed, catalytically competent conformation of PimA. These observations raise the possibility that ML0451c not only scaffolds the complex but also regulates PimA activity through conformational control.

EphE belongs to the α/β-hydrolase superfamily, a large and evolutionarily ancient family of enzymes that catalyze diverse hydrolytic reactions using a catalytic triad^38^. Although EphE is the only annotated epoxide hydrolase whose depletion causes a severe growth defect in *M. tuberculosis*^39^, its biochemical function remains unclear. Unlike canonical epoxide hydrolases, EphE lacks the conserved nucleophilic Asp or Ser residue located at the highly conserved “nucleophile elbow” and recombinant EphE exhibits very little epoxide hydrolase activity on a 9,10-cis-epoxystearic acid substrate *in vitro*^40^. The predicted active site retains residues corresponding to the catalytic acid–base pair (Asp296 and His297) but contains Ala122 in place of the canonical nucleophile (Figure 3D, inset 2). Together, these features suggest that EphE has evolved a distinct enzymatic function from other members of the epoxide hydrolase family.

One intriguing possibility is that EphE functions as the long-sought PIP phosphatase that converts PIP to PI prior to PimA-dependent mannosylation. In this case, EphE would employ a non-canonical catalytic mechanism, involving its His-Asp catalytic dyad rather than the Asp/Ser-centered architecture characteristic of classical epoxide hydrolases. Such a function would provide a parsimonious explanation for EphE’s near-essentiality, its lack of detectable epoxide hydrolase activity, its predicted association with PimA, and the severe PIM deficiency observed in the *ΔephE* mutant. Additional support for this model comes from analysis of the electrostatic surface properties of the predicted complex. The putative EphE active site contains several conserved polar and positively charged residues, including His121, Arg199, and Gln243, that form a solvent-accessible cavity adjacent to the membrane interface (Figure 3D, inset 2). This cavity is embedded within a strongly electropositive surface patch that faces the inner membrane, positioning it to interact with negatively charged phospholipid headgroups (Figure S3A-B). The uniquely positive electrostatic distribution of EphE (Figure S3C-D) may be required for it to interact with the highly anionic headgroups of phosphorylated membrane lipids such as PIP. Although direct biochemical evidence for PIP phosphatase activity is currently lacking, the structural prediction, genetic, and TLC data collectively identify EphE as a strong candidate for this missing step in mycobacterial PIM biosynthesis.

### ML2271 and ML0614 are essential cofactors of the MptB mannosyltransferase

MptB (ML0591c) is an essential GT-C-family mannosyltransferase that is involved in the synthesis of Ac_1_/Ac_2_PIM_4_, by using a lipid linked mannose donor (polyprenol-P-mannose) to add mannose residues to Ac_1_/Ac_2_PIM_2_ and/or Ac_1_/Ac_2_PIM_3_^16^. MptB functions together with LpqW, a periplasmic lipoprotein that channels the PIM_4_ products of MptB into the LM and LAM pathways^41–43^. Consistent with their established functional relationship, *mptB* and *lpqW* knockdown or knockout strains exhibited highly similar chemical-genetic profiles in all three species analyzed (Figure S4A-C), driven primarily by hypersensitivity to diverse peptidoglycan-targeting antibiotics^10–12^. In addition to LpqW, AlphaFold3 additionally predicts interactions between MptB and two previously uncharacterized essential proteins: ML2271 and ML0614 (size-corrected ipTM = 0.58 and 0.56, respectively (Figure 4A). The phenotypes of *ML2271* homologs were strongly correlated with those of *mptB*-pathway genes, including *lpqW* and *mptA*, across all three mycobacterial species, whereas *ML0614* displayed weaker phenotypic similarity (Figure S4A-C). *ML2271* encodes a DUF3592-family membrane protein of unknown function, while *ML0614* is a small 95-amino-acid protein predicted to contain a single transmembrane helix and a short periplasmic helix.

**Figure 4.**
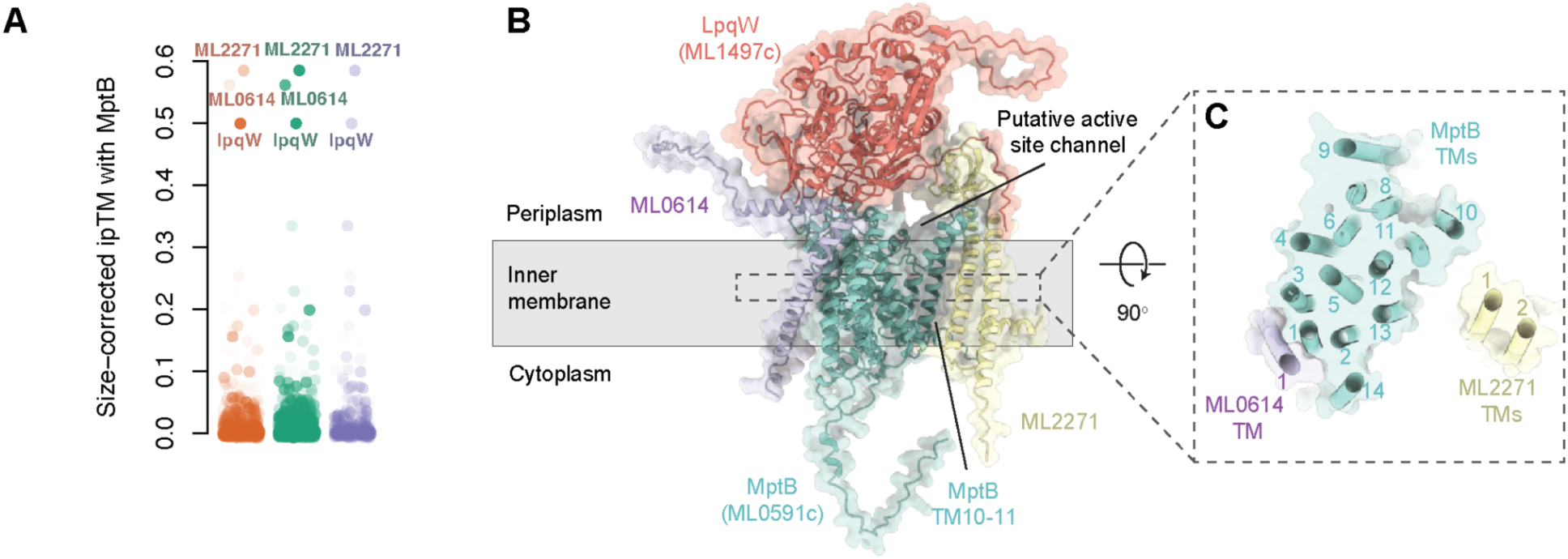
MptB–LpqW–ML0614–ML2271 is predicted to form a membrane-associated complex. **(A)** Size-corrected ipTMs between MptB (ML0591c/MSMEG3120/Rv1459c/Cg1766) and all other proteins in the *M. leprae* data. Opacity of points represents the phenotypic correlations in *M. smegmatis* (orange), *M. tuberculosis* (green), and *C. glutamicum* (purple). ML1497c (LpqW/MSMEG5130/Rv1166/Cg1249), ML0614 (MSMEG4534/Rv2401A), and ML2271 (MSMEG1110/Rv0556/Cg0553) (labeled) have size-corrected ipTM > 0.45, and are phenotypically correlated in all datasets. **(B)** AlphaFold3 model of the *M. leprae* MptB-associated complex. MptB (ML0591c, cyan), LpqW (ML1497c, salmon), ML0614 (purple), and ML2271 (yellow) are shown as ribbon representations overlaid with transparent molecular surfaces and positioned relative to the inner membrane (grey). LpqW forms an extensive periplasmic domain above the membrane, whereas MptB comprises a 14-transmembrane-helix domain. ML0614 and ML2271 associate with the periphery of the complex through single- and two-pass transmembrane segments, respectively. A putative active-site channel extending from the membrane toward the periplasmic region is indicated. **(C)** Cross-sectional view of the transmembrane region following a 90° rotation relative to (B). Transmembrane helices of MptB, ML0614 and ML2271 are shown as cylinders and numbered. The transmembrane segments of ML0614 and ML2271 are positioned adjacent to the MptB transmembrane domain with ML2271 flanking the putative active-site channel of MptB.

The predicted structure of the MptB–LpqW–ML0614–ML2271 complex provides insight into the potential functions of these novel partners (Figure 4B). MptB consists of a transmembrane domain composed of 14 transmembrane helices (TMs), where TM10 and TM11 are notably splayed away from the main transmembrane bundle (Figure 4B), creating a periplasmic-facing cavity that houses its putative active site. As expected, LpqW associates extensively with the periplasmic face of MptB and contacts regions adjacent to this catalytic center, consistent with its experimentally established role in committing the product of MptB to the LM/LAM pathway^41–43^. Remarkably, ML2271 contains two transmembrane helices connected by a small β-barrel domain and directly flanks the splayed out TM10 and TM11, placing it in a strategic position to potentially influence substrate access to, or product release from, the MptB active site (Figure 4C). The β-barrel domain of ML2271 contacts the N-terminus LpqW, collectively forming a channel that connects the active site of MptB to the periplasm (Figure 4B). Such channels have been proposed to facilitate substrate movement and processive glycan elongation in other membrane-associated glycosyltransferases^44^. Intriguingly, a structurally analogous channel is also observed in the predicted MptA–LmeA complex, suggesting that formation of an auxiliary substrate-guiding channel may represent a conserved architectural feature of processive mannosyltransferases involved in LM/LAM biosynthesis. In contrast, ML0614 binds adjacent to TM1, TM2, and TM14 of MptB (Figure 4C) and its short periplasmic helix create an interaction surface for MptB and LpqW to bind (Figure 4B), suggesting a role in stabilizing the complex.

Taken together, the genetic and structural data strongly support ML2271 as a previously unrecognized essential cofactor of MptB, likely contributing to substrate handling or processive glycan extension through formation of a channel adjacent to the catalytic site. ML0614 also appears to be a bona-fide component of the MptB complex, although its role is less clear and may involve stabilization of the MptB-LpqW complex.

### AftD may license mycolic acid export through a direct interaction with TmaT

Mycolic acids are the defining components of the mycomembrane. These essential long-chain fatty acids are esterified to the terminal arabinan residues of AG, forming the inner leaflet of the mycomembrane. Because mycolic acids are covalently attached to AG, the synthesis and export of these two components is likely coordinated. Our predicted PPI data suggest that a physical interaction between AftD (ML2570) and TmaT (ML2508) may coordinate these processes. AftD is essential arabinosyltransferase involved in the biosynthesis of the arabinan domains of AG and LAM^45,46^ and TmaT is an essential transmembrane acyltransferase-superfamily (TmAT)^47^ enzyme that acetylates trehalose monomycolate and enables subsequent mycolic acid transport by MmpL3^48,49^ (Table S2). AftD and TmaT have both a strong predicted interaction (size-corrected ipTM = 0.71; Figure 5A) and some of the strongest phenotypic correlations observed across all three species examined (*M. smegmatis*, *r* = 0.90; *M. tuberculosis*, *r* = 0.96; *C. glutamicum*, *r* = 0.95; Figure S5), suggesting a physical and functional interaction. Second, and critically, even though *aftD* has a well-established role in arabian assembly, *aftD* knockdowns/knockouts phenotypically cluster with genes involved in mycolic acid synthesis and transport rather than with other AG biosynthetic genes (Figure S5A-C), suggesting that AftD specifically affects mycolic acid biogenesis in addition to arabinan assembly. This hypothesis is consistent with previous reports showing that AftD physically interacts with MmpL3 and PgfA (a MmpL3-interacting protein involved in trehalose monomycolate export)^50,51^.

**Figure 5.**
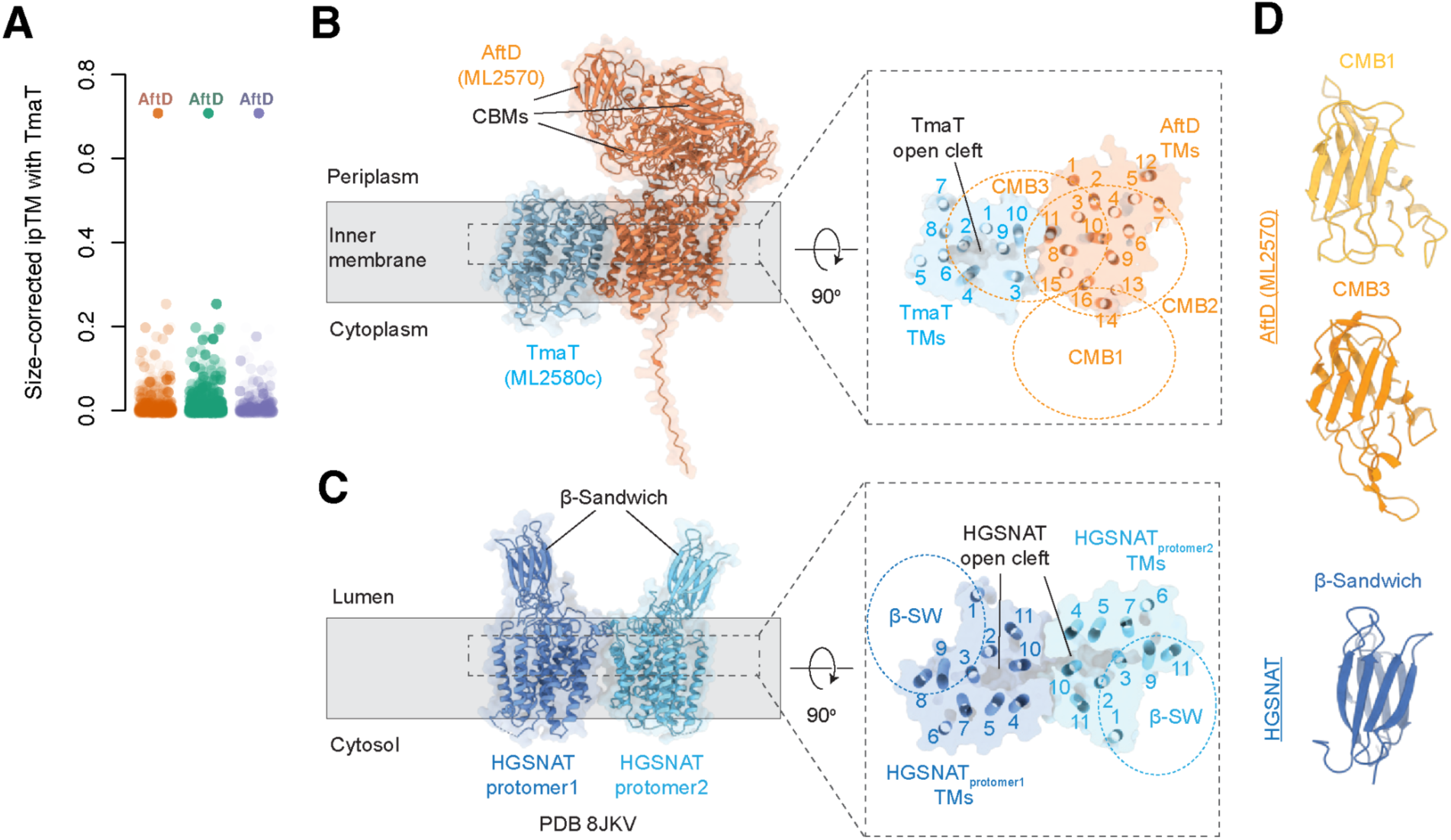
Structural parallels between predicted AftD-TmaT complex and HGSNAT. **(A)** Size-corrected ipTMs between TmaT (ML2580c/MSMEG0319/Rv0228/Cg3163) and all other proteins in the *M. leprae* data. Opacity of points represents the phenotypic correlations in *M. smegmatis* (orange), *M. tuberculosis* (green), and *C. glutamicum* (purple). AftD (ML2570/MSMEG0359/Rv0236c/Cg3161) exhibits both a strong predicted interaction and highly correlated phenotypes in all three datasets. **(B)** AlphaFold3 model of the *M. leprae* AftD–TmaT complex. AftD (ML2570, orange) and TmaT (ML2580c, cyan) are shown as ribbon representations overlaid with transparent molecular surfaces and positioned relative to the inner membrane (grey). The periplasmic carbohydrate binding modules (CBMs) of AftD are indicated. Inset: Cross-sectional view of the AftD–TmaT transmembrane region following a 90° rotation relative to panel **A**. Transmembrane helices are shown as cylinders and numbered. AftD comprises 16 transmembrane helices, whereas TmaT contains 10 transmembrane helices that pack against the AftD transmembrane domain. An open cleft is formed at the AftD–TmaT interface. The positions of the periplasmic CBMs are indicated by the dotted circles. **(C)** Dimeric structure of the human lysosomal acetyltransferase HGSNAT (PDB: 8JKV^52^) shown in the same orientation as in panel **A**. Individual protomers are coloured dark blue and light blue. The luminal β-sandwich (β-SW) domain of each HGSNAT protomer is indicated. Inset: Equivalent cross-sectional view of the HGSNAT dimer following a 90° rotation relative to panel **B inset**. Transmembrane helices from each protomer are shown as cylinders and numbered. The dimer interface forms an open cleft analogous to that observed in the AftD–TmaT complex. The position of the luminal β-SW domain within each HGSNAT protomer is indicated. **(D)** Comparison of the CBMs AftD and HGSNAT. AftD contains three CBMs, two of which (CBM1 and CBM3) are oriented toward TmaT, whereas HGSNAT contains a single β-sandwich domain. The AftD CBMs contain β-sandwich folds that are analogous to the HGSNAT β-SW domain.

To better understand how AftD may affect TmaT activity, we examined the predicted heterodimer of these two proteins and compared it to that of the homodimeric human lysosomal acetyltransferase (HGSNAT)^52^, the only structurally characterized member of the TmAT superfamily (Figure 5B-C). AftD and TmaT interact through their transmembrane domains, with the 10 transmembrane helices of TmaT packing against the 16-transmembrane-helix core of AftD (Figure 5B inset). In particular, TM11 of AftD contacts TM10 of TmaT, while TM15 of AftD interacts with TmaT TM3 (Figure 5B inset). This interaction creates a periplasmic-facing open cleft in TmaT that sits below the periplasmic domain of AftD, which harbors three carbohydrate binding modules (denoted as CBM1 (residues 704-844), CBM2 (residues 857-1170), and CBM3 (residues 945-1092) (Figure 5B inset). Strikingly, the overall architecture of the AftD–TmaT complex resembles that of HGSNAT. In HGSNAT, two protomers associate through their transmembrane domains to form an obligate dimer, creating an open lumenal cleft analogous to that observed in the AftD–TmaT complex (Figure 5C inset).

The structural similarity between the AftD-TmaT complex and the HGSNAT dimer suggests several mechanisms through which AftD could regulate TmaT activity. First, in HGSNAT, the lumenal loops connecting TM4-TM5 and TM10-TM11 contribute substantially to the dimerization interface, and this dimerization is required for enzymatic activity^52^. AftD may function as a structural surrogate for the second HGSNAT protomer, interacting with the equivalent parts of TmaT to stabilize a functional conformation. Second, AftD may promote substrate access to the catalytic site of TmaT. In HGSNAT, TM10 forms part of the solvent-accessible tunnel that accommodates Ac-CoA entry from the cytosolic side of the membrane^52^ (Figure 5C inset). AftD is predicted to interact directly with TmaT TM3 and TM10, placing AftD in an ideal position to influence the conformation of the open cleft observed in TmaT (Figure 5B inset). We therefore speculate that AftD may stabilize TmaT in an open, catalytically competent state. Third, AftD may contribute directly to TmaT substrate recognition. Many TmAT-family enzymes possess accessory substrate-binding domains that help position carbohydrate acceptors (e.g., the lumenal β-sandwich domain (β-SW) of HGSNAT^52^, or the SGNH domain of many bacterial TmaTs^53^). Although TmaT lacks an obvious substrate-binding domain, AftD contains three CBMs that may play such a role, particularly CBM1 and CBM3, which is flexible^45^, are oriented toward the open cleft of TmaT. Intriguingly, the AftD CBMs structurally resemble the β-sandwich domain of HGSNAT (Figure 5D), suggesting a potential role in substrate binding or catalysis. Together, these data suggest that the interaction between AftD and TmaT is required to license mycolic acid export and suggest a mechanism by which this may occur.

### ML1347 activates MtrP to regulate mycolic acid export

MtrP is an essential S-adenosylmethionine (SAM)-dependent methyltransferase encoded within a conserved genetic locus that includes *tmaT*^48^, *pgfA*^51^, and other genes implicated in mycolic acid transport and polar envelope biogenesis^54^. Although previous work established a role for MtrP in regulating mycolic acid export^54^, the molecular basis of its activity has remained unclear. Our data suggest that MtrP functions together with ML1347, a small but essential UPF0434-family protein that exhibits homology to the eukaryotic methyltransferase activator Trm112. Consistent with this model, ML1347 and MtrP have a strong predicted interaction (Figure 6A), and knockdown of the *ML1347* homolog in *M. tuberculosis* (*rv1684*) phenocopied the knockdown/knockout of *mtrP* (r = 0.92) (Figure 6B). We were unable to evaluate this relationship in *M. smegmatis*, where *mtrP* (*MSMEG0310*) is annotated as a pseudogene and was therefore not targeted in the CRISPRi screen, or in *C. glutamicum*, where the ML1347 homolog (*cg1592*) was absent from the available Tn-seq data due to its essentiality.

**Figure 6.**
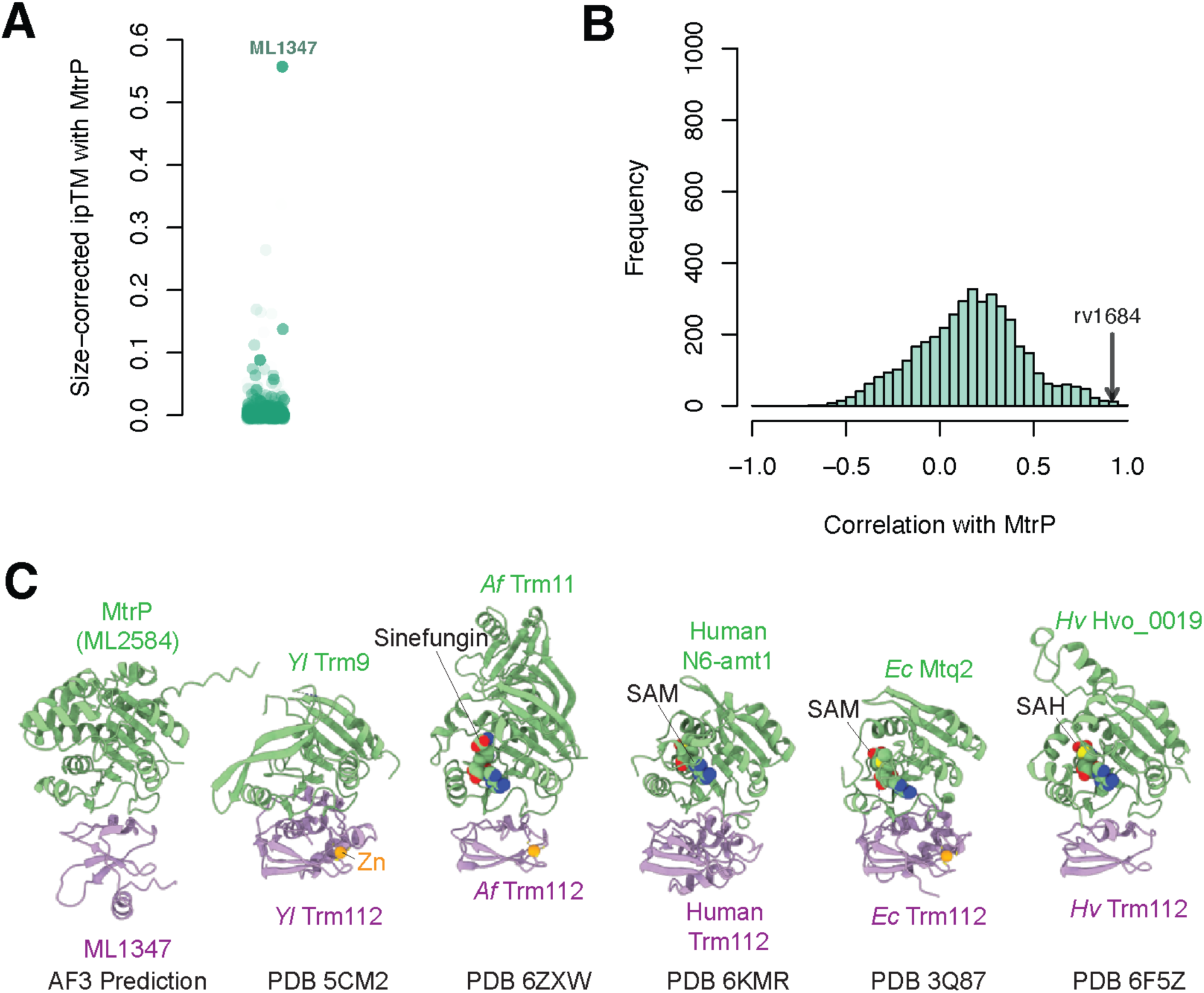
ML1347 may function as a co-factor of MtrP. **(A)** Size-corrected ipTMs between MtrP (ML2584/Rv0224c) and all other proteins in the *M. leprae* data. Opacity of points represents the phenotypic correlations in *M. tuberculosis* (green). **(B)** Histogram of phenotypic correlations between *mtrP* and all other genes in *M. tuberculosis*, showing strong phenotypic correlation to the *ML1347* homolog (*rv1684,* black arrow). **(C)** AlphaFold3 model of the *M. leprae* MtrP–ML1347 complex compared with representative structures of Trm112-associated methyltransferases from archaea and eukaryotes. MtrP (ML2584, green) and ML1347 (purple) are shown alongside the *Yarrowia lipolytica* (*Yl*) Trm9–Trm112 complex (PDB: 5CM2^56^), *Archaeoglobus fulgidus* (*Af*) Trm11–Trm112 complex (PDB: 6ZXW^57^), human N6AMT1–TRMT112 (PDB: 6KMR^58^), *Encephalitozoon cuniculi* (*Ec*) Mtq2–Trm112 complex (PDB: 3Q87^59^), and *Haloferax volcanii* Hvo_0019–Trm112 complex (PDB: 6F5Z^60^). Bound cofactors and ligands, including S-adenosylmethionine (SAM), S-adenosylhomocysteine (SAH), sinefungin, and a structural Zn²⁺ ion, are shown as spheres. MtrP adopts the characteristic class I S-adenosylmethionine-dependent methyltransferase fold and forms a heterodimeric complex with ML1347 analogous to the conserved Trm112-associated methyltransferase family.

In eukaryotes, Trm112 associates with several essential SAM-dependent methyltransferases and is required for their activity^55^. To determine if the predicted ML1347-MtrP interaction resembles that of its eukaryotic homologs, we compared the structure of the predicted MtrP–ML1347 complex with the experimental structures of known Trm112-associated methyltransferases, including eukaryotic Trm9–Trm112^56^, Trm11–Trm112^57^, N6-amt1–Trm112^58^, Mtq2–Trm112^59^, and the archaeal Hvo_0019–Trm112 complexes^60^ (Figure 6C). We found that MtrP adopts the canonical class I SAM-dependent methyltransferase fold and forms a heterodimeric complex with ML1347 that closely resembles known Trm112-associated methyltransferases, suggesting that ML1347 performs a conserved cofactor-like role despite limited primary sequence similarity with Trm112.

Our predicted protein–protein interaction network additionally identified an interaction between ML1347 and MenG (ML2273), the demethylmenaquinone methyltransferase responsible for the final methylation step in menaquinone biosynthesis^61^. Although not supported by our phenotypic data, this interaction would be consistent with the multifunctional nature of Trm112 in eukaryotes, with pulldown data associating the Trm112 homolog with MenG in the archaeon *Haloferax volcanii*^60^, and with the prominent hydrophobic surface patch on MenG that resembles the ML1347-binding surface of MtrP, suggesting that ML1347 may also separately activate MenG (Figure S6A-B). Taken together, our combined phenotypic and predicted PPI data suggest that ML1347 is required to activate MtrP, thereby enabling the methylation-dependent processes that regulate mycolic acid export and may also activate MenG. These findings extend the Trm112 methyltransferase activator paradigm beyond archaea and eukaryotes and identify a previously unrecognized bacterial methyltransferase regulatory module linked to cell envelope biogenesis.

### Ferritin-superfamily proteins ML1560-ML1561 activate the essential DD-carboxypeptidase DacB (ML0211)

Peptidoglycan (PG), an essential glycopolymer conserved in almost all bacteria, undergoes continuous remodeling to maintain cell shape, coordinate cell division, and withstand environmental stresses. DD-carboxypeptidases (DD-CPases) play an important role in PG remodeling by hydrolyzing the terminal D-alanine residues from pentapeptide stem peptides to modulate the extent of PG cross-linking. DD-CPases are members of the S11, S12, and S13 peptidase families. *M. leprae, M. tuberculosis, M. smegmatis, and C. glutamicum* each possess a single S13-family DD-CPase (ML0211, Rv3627c, MSMEG6113, and Cg2987, respectively), together with one or more S11-family DD-CPases. In both *M. tuberculosis* and *M. smegmatis*, the S13-family enzyme is essential, whereas the S11-family enzymes are dispensable, indicating that the S13-family DD-CPases are the major drivers of PG remodeling in these bacteria.

In our predicted PPI data, we identify a previously unrecognized functional association between the *M. leprae* S13-family DD-CPase DacB (ML0211) and two DUF4439-family proteins, ML1560 and ML1561. In the co-folded model, ML0211 interacts directly with ML1561 (size-corrected ipTM = 0.62), which in turn associates with ML1560 (size-corrected ipTM = 0.61), forming a ternary complex (Figure 7A). ML1560 and ML1561 bind one face of DacB, where ML1561 forms an extensive protein–protein interaction interface adjacent to the catalytic domain. Notably, the conserved catalytic SXXK, SXN, and KTG motifs remain fully solvent accessible (Figure 7B-C), indicating that ML1560 and ML1561 do not occlude the active-site cleft but instead are positioned to potentially regulate DacB through structural stabilization or allosteric modulation.

**Figure 7.**
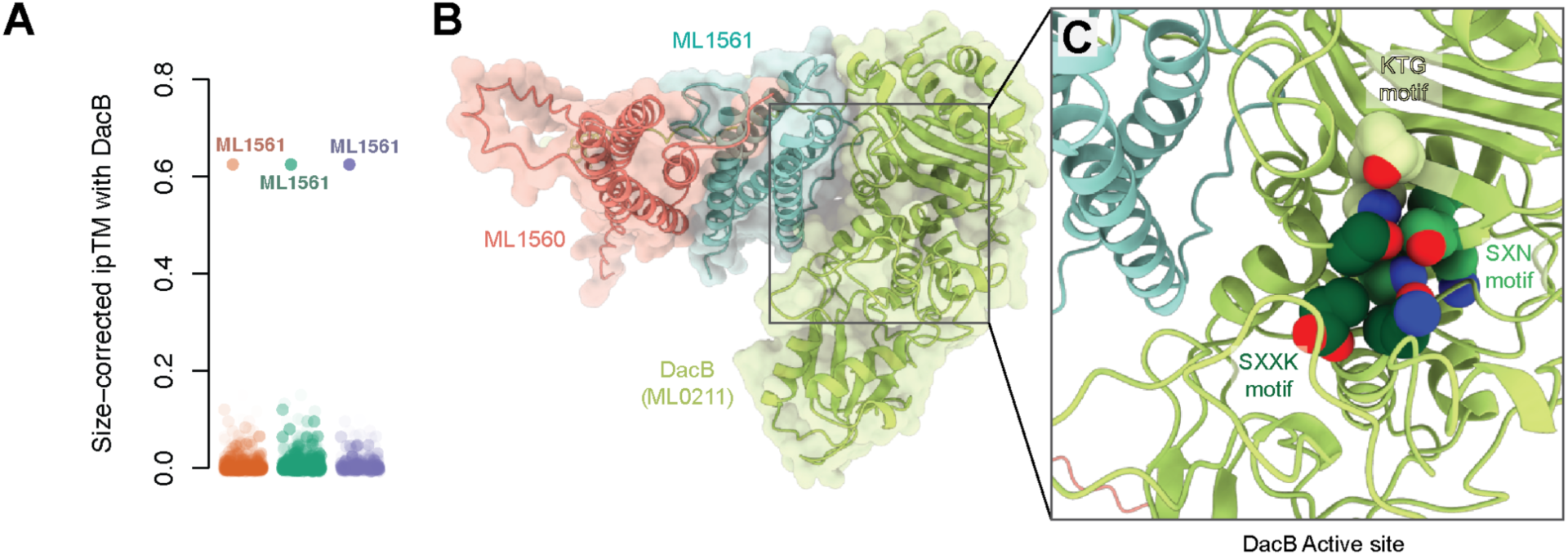
Predicted structure of the DacB–ML1560–ML1561 complex. **(A)** Size-corrected ipTMs between DacB (ML0211/MSMEG6113/Rv3627c/Cg2987) and all other proteins in the *M. leprae* data. Opacity of points represents the phenotypic correlations in *M. smegmatis* (orange), *M. tuberculosis* (green), and *C. glutamicum* (purple). ML1561 (MSMEG2622/Rv2844/Cg2180) exhibits high size-corrected ipTM and phenotypic correlations in all three datasets. **(B)** AlphaFold3 model of the *M. leprae* DacB-associated complex. DacB (ML0211, green), ML1561 (cyan), and ML1560 (salmon) are shown as ribbon representations overlaid with transparent molecular surfaces. ML1560 and ML1561 associate with one face of DacB adjacent to the catalytic domain, forming an extended protein–protein interaction interface. **(C)** Close-up view of the DacB active site. Conserved catalytic motifs characteristic of class B penicillin-binding proteins, including the SXXK, SXN and KTG motifs, are indicated and shown as spheres. The catalytic pocket remains solvent-accessible in the predicted complex, with ML1560 and ML1561 positioned adjacent to, but not occluding, the active-site cleft.

This predicted interaction is strongly supported by comparative chemical genomics. Across all three datasets, homologs of ML0211 consistently co-cluster with DUF4439-family proteins (represented by the fused protein Cg2180 in *C. glutamicum*) (Figure S7A-C), driven by shared resistance to vancomycin and, in *M. smegmatis* and *C. glutamicum*, increased sensitivity to specific cephalosporins. These conserved antibiotic susceptibility profiles suggest that the S13-family DD-CPase and DUF4439 proteins participate in a common pathway governing peptidoglycan homeostasis. However, since DUF4439-family proteins in *M. smegmatis* and *M. tuberculosis* are not essential, this requirement does not appear to be absolute and may depend on stress conditions.

Although DUF4439 proteins adopt a ferritin-like fold, their function appears distinct from the canonical role of ferritins in intracellular iron storage. Instead, the *M. tuberculosis* homolog of ML1560 (Rv2843) has been experimentally identified as a substrate of the twin-arginine translocation (Tat) pathway^62^, placing it within the cell envelope where interaction with the periplasmic DD-carboxypeptidase is physically plausible. This observation is further supported by studies in the distantly related Firmicute *Listeria monocytogenes*, where the sole ferritin-like protein localizes to the cell wall^63^ and contributes to envelope integrity during β-lactam stress^64^, suggesting that ferritin-like proteins may have evolved conserved roles in regulating cell envelope biogenesis beyond iron metabolism.

### Protein mannosylation is carried out by a heteropentameric complex

O-mannosylation is a post-translational modification of secreted proteins broadly conserved across eukaryotes and Actinobacteria and essential for *Mycobacterium tuberculosis* virulence^65^. In eukaryotes, protein mannosylation is catalyzed by homo- or heterodimeric GT39-family protein O-mannosyltransferases that utilize a lipid-linked mannose donor (e.g., dolichol-phosphate-mannose). Mycobacteria encode a single GT39-family enzyme, Pmt, which is absolutely required for protein mannosylation^66^. Unlike eukaryotic PMTs, which contain large lumenal MIR domains and function as obligate dimers^67^, actinobacterial Pmt proteins lack substantial periplasmic domains and dimerization has not been observed.

Our pairwise PPI predictions identified three conserved proteins predicted to interact with the *M. leprae* Pmt homolog ML0192c: ML0810c, ML1065, and ML2687c (size-corrected ipTMs of 0.66, 0.51, and 0.45, respectively; Figure 8A). Knockdown/knockout phenotypes of all three genes were strongly correlated to those of *pmt* in all three bacteria (Figure S8A-C), albeit their phenotypes were generally of lower magnitude than those of *pmt* (Figure S8D). To determine whether these proteins form a stable assembly, we introduced a C-terminal 3×FLAG tag at the endogenous *M. smegmatis pmt* locus (*MSMEG5447*) and performed affinity purification followed by mass spectrometry (Figure 8B; Methods). Th*e M. smegmatis* homologs of our predicted interaction partners [MSMEG6899 (ML2687c homolog), MSMEG5083 (ML1065 homolog), and MSMEG1929 (ML0810c homolog)] were the most abundant additional proteins recovered in the MSMEG5447 pulldown (Table S5), providing strong experimental support for the existence of this complex *in vivo*.

**Figure 8.**
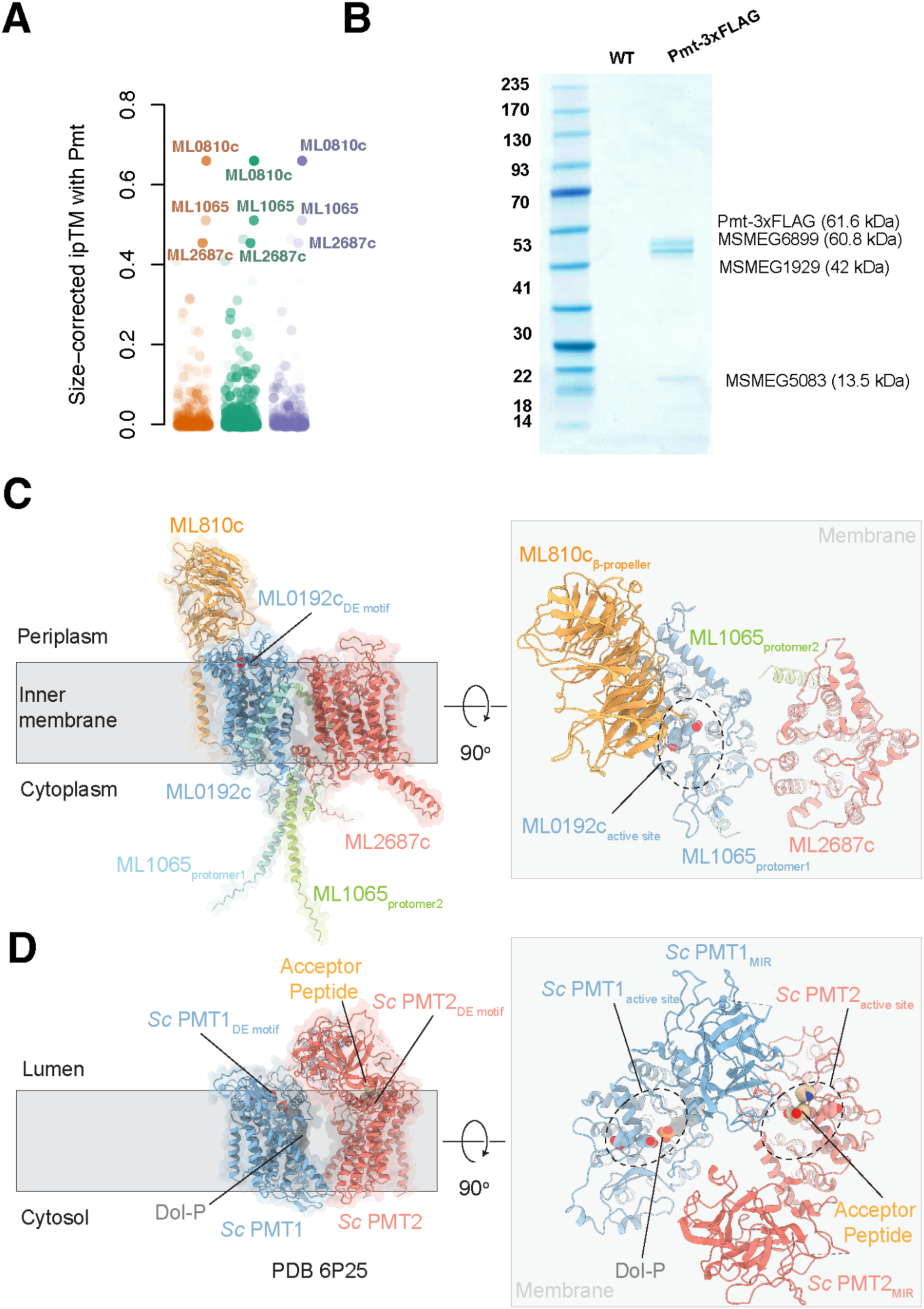
Protein mannosylation is performed by a heteromeric protein complex. **(A)** Size-corrected ipTMs between Pmt (ML0192c/MSMEG5447/Rv1002c/Cg1014) and all other proteins in the *M. leprae* data. Opacity of points represents the phenotypic correlations in *M. smegmatis* (orange), *M. tuberculosis* (green), and *C. glutamicum* (purple). ML0810c (MSMEG1929/Rv3212/Cg0880), ML1065 (MSMEG5083/Rv1209/Cg1264), and ML2687c (MSMEG6899/Rv0051/Cg3312) exhibit high size-corrected ipTM and phenotypic correlations in all three datasets. **(B)** Pmt-3xFLAG pulldown. *M. smegmatis* membranes were solubilized and purified using α-FLAG resin. The elution fractions of wildtype (WT) and Pmt-3xFLAG cells were analyzed with SDS-PAGE. Bands and their molecular weights of protein corresponding to Pmt components are indicated. **(C)** AlphaFold3 model of the *M. leprae* PMT complex. ML0192c (blue), ML1065 (light green and cyan), ML2687c (salmon), and ML810c (orange) are shown as ribbon representations overlaid with transparent molecular surfaces and positioned relative to the inner membrane (grey). The conserved catalytic Asp–Glu (DE) motif of ML0192c is indicated. ML810c forms a periplasmic β-propeller domain associated with the membrane-embedded core of the complex. Inset: Periplasmic view of the *M. leprae* PMT complex following a 90° rotation. ML0192c and ML2687c form the central catalytic core, with the active site of ML0192c positioned beneath the β-propeller domain of ML810c. ML1065 associates with the periphery of the complex and forms a dimeric scaffold that bridges ML0192c and ML2687c. **(D)** Structure of the *Saccharomyces cerevisiae* (*Sc*) PMT1–PMT2 heterodimer (PDB: 6P25^67^) shown in the same orientation as in the putative mycobacterial Pmt complex in (C). PMT1 and PMT2 are colored blue and salmon, respectively. The catalytic DE motifs, dolichyl phosphate (Dol-P), and acceptor peptide are indicated. Inset: Equivalent view of the Sc PMT1–PMT2 complex following a 90° rotation. The active sites of PMT1 and PMT2, bound Dol-P, acceptor peptide, and luminal MIR domains are indicated. Comparison with the *M. leprae* complex highlights similarities in the spatial organization of the catalytic center relative to the membrane and luminal domains despite differences in subunit composition.

To determine the possible roles of these accessory proteins, we co-folded all four proteins using AlphaFold3 (Figure 8C) and compared the structure of the predicted complex to the eukaryotic Pmt1-Pmt2 heterodimer (Figure 8D). The resulting model revealed a highly organized membrane-associated complex whose overall architecture bears striking resemblance to the eukaryotic Pmt1–Pmt2 heterodimer^67^ despite the absence of obvious sequence homology outside the GT39 catalytic core.

ML0192c (Pmt) formed a heterodimer with ML2687c, a GT87-family glycosyltransferase-like protein that lacks the conserved catalytic residues required for enzymatic activity^68^. This arrangement is reminiscent of the yeast Pmt1–Pmt2 complex^67^, in which Pmt1 catalyzes mannosylation, while Pmt2 recruits the acceptor polypeptide (Figure 8D 90° rotation), suggesting that ML2687c may play a non-catalytic role similar to Pmt2 (Figure 8C 90° rotation). This is consistent with the observation that inactivation of ML2687c homologs significantly affects complex activity (95%, 39%, and 11% of the *pmt* phenotype in *M. tuberculosis*, *M. smegmatis*, and *C. glutamicum* respectively; Figure S8D), despite the likely lack of catalytic activity. This dimeric organization appears to be shared not only with eukaryotic Pmts, but also with distantly related GT39-family enzymes from Paenibacillus species, which are translationally fused to a GT87-family enzyme (i.e., UniProt V9WAU2), suggesting that dimerization may be a general property of Pmts.

ML0192c and ML2687c are predicted to interact only modestly with one another at the cytoplasmic and periplasmic regions, leaving a cavity between the two proteins in the transmembrane region similar to (but smaller than) that observed in the Pmt1–Pmt2 heterodimer^67^. In our predicted complex, the interaction between ML0192c and ML2687c appears to be stabilized by the bridging of two protomers of ML1065, a small DivIVA-family protein containing two α-helical domains separated by a flexible linker (Figure 8C). The N-terminal helical regions of ML1065 mediate dimerization, whereas the C-terminal helices contact the transmembrane regions of ML0192c and ML2687c on opposite sides of the assembly (Figure 8C 90° rotation). This arrangement effectively links the two GT-C proteins into a single functional unit and may additionally couple the complex to membrane-localized complexes (such as the translocon or translating ribosomes). However, the role of ML1065 appears largely dispensable: inactivation of ML1065 homologs minimally phenocopied the *pmt* phenotype (0%, 57%, and 25% of the *pmt* phenotype in *M. smegmatis*, *M. tuberculosis*, and *C. glutamicum*, respectively; Figure S8D).

The fourth component, ML0810c, forms a large periplasmic eight-bladed β-propeller domain positioned directly above the catalytic center of ML0192c (Figure 8C 90° rotation). The active site of ML0192c lies beneath this β-propeller, suggesting that ML0810c potentially functions as a gatekeeper for substrate access. This organization closely parallels the role of the lumenal MIR domains in the eukaryotic Pmt1–Pmt2 complex, which cap the catalytic center (Figure 8D 90° rotation) and contribute to acceptor-protein recognition^69^. Although ML0810c is structurally unrelated to MIR domains, its analogous position above the active site strongly suggests a similar role in substrate selection. This model is supported by the importance of ML0810 homologs for complex activity: their inactivation phenocopies *pmt* in *M. tuberculosis* and *C. glutamicum* and causes 62% of the *pmt* phenotype in *M. smegmatis* (Figure S8D).

Although the exact role of protein O-mannosylation in actinobacteria remains enigmatic, it appears to be important for the stabilization and/or function of certain periplasmic proteins, including PG degrading enzymes (e.g. Ami3^25^). Our identification of the additional members of this complex opens the door to a deeper understanding of the mechanisms and regulation of this important post-translational modification.

### ML1026 and ML1638 are novel RNA-polymerase interacting proteins

RNA polymerase (RNAP) is the central enzyme of the bacterial transcription machinery and the direct target of rifampicin, a cornerstone of first-line tuberculosis therapy^17^. Mycobacterial RNAP has been extensively studied, leading to the discovery of several lineage-specific transcription factors and accessory subunits, including CarD^70^ and RbpA^71^, which are critical regulators of transcription initiation. Consistent with these established roles and published structures^71,72^ both CarD and RbpA exhibited strong predicted interactions with the RNAP core subunits, RpoB (β) and RpoC (β′) (size-corrected ipTM CarD-RpoB = 0.45; size-corrected ipTM RbpA-RpoC = 0.65; Table S2). To identify additional RNAP-associated proteins, we plotted the size-corrected ipTM scores for all *M. leprae* proteins against RpoB and RpoC (Figure 9A). This analysis identified two previously unrecognized RNAP-binding proteins: ML1026, an overlooked but essential protein, and ML1638, a representative of a broadly distributed yet poorly characterized family of RNAP-associated factors.

**Figure 9.**
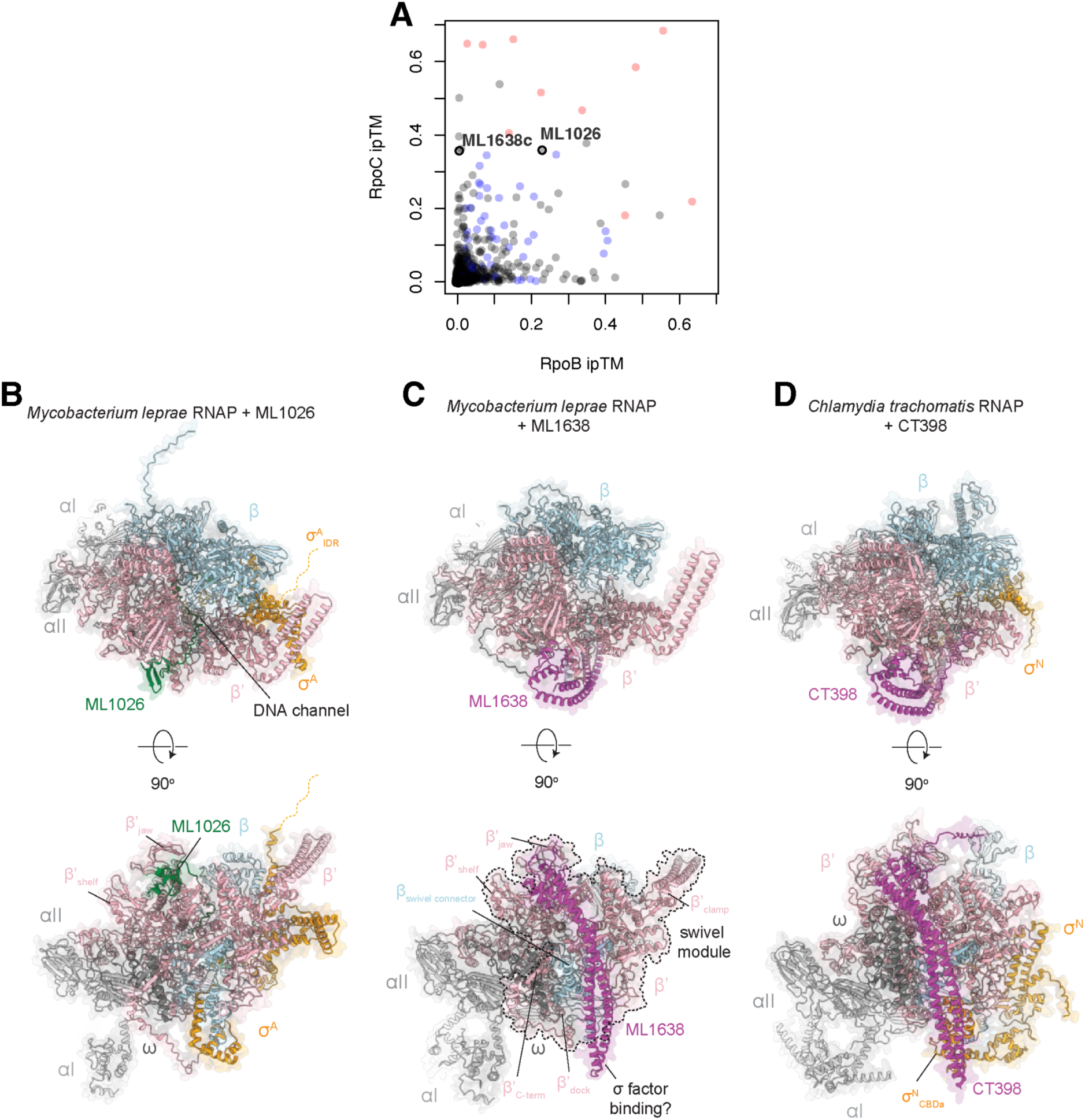
Pooled-AF3 identifies new RNA-polymerase interacting proteins. **(A)** Plot of size-corrected ipTMs for RpoB and RpoC, highlighting known RNA-polymerase interacting proteins (in red), weaker predicted interactions with ribosomal proteins (blue), and new RNA-polymerase interacting proteins ML1026 and ML1638. **(B)** AlphaFold3 model of the *M. leprae* RNA polymerase (RNAP) holoenzyme in complex with ML1026. RNAP subunits are shown as cartoon ribbon representations overlaid with transparent molecular surfaces and coloured individually, with ML1026 shown in green. Orthogonal views reveal association of ML1026 with the β′ jaw and β′ shelf regions on the exterior surface of RNAP, with its N-terminus buried in the downstream DNA channel and active site formed by β and β’. The intrinsically disordered region (IDR) of σ^A^ is omitted for clarity and indicated by an orange dashed line. **(C)** AlphaFold3 model of the *M. leprae* RNAP core enzyme in complex with ML1638 (magenta). Orthogonal views show that ML1638 occupies a binding site adjacent to the β′ shelf and β′ jaw domains and extends across the RNAP swivel module. The distal tip of the coiled-coil overlaps with the canonical σ-factor binding site on RNAP. **(D)** Predicted structure of the *Chlamydia trachomatis* σ^N^-holoenzyme in complex with the ML1638 homolog CT398, shown in the same orientation as in (C) and (D). CT398 (magenta) associates with the β′ shelf and β′ jaw domains and extends across β′ through an elongated coiled-coil similar to ML1638. The distal tip of the coiled-coil is predicted to interact with the core-binding domain (CBDa) of σ^N^.

ML1026 is a small DUF4193-family protein that is widely conserved throughout Actinobacteria and is essential for growth in *M. tuberculosis*^39^. ML1026 exhibited two intermediate interactions, both with components of RNAP (Table S2; size-corrected ipTM with RpoB = 0.23; size-corrected ipTM with RpoC = 0.36). Notably, depletion of the *M. tuberculosis* homolog (*rv2699c*) sensitizes cells to rifampicin^12^, providing independent evidence for a functional relationship between ML1026 and RNAP. Co-folding ML1026 with the RNAP σ^A^-holoenzyme (α₂ββ′ωσ^A^) suggested that it binds the β and β’ subunits with high ipTM to both subunits (raw ipTMs: 0.7-0.9). ML1026 consists of a C-terminal globular domain that binds adjacent to the β′ jaw and β′ shelf domains of RNAP, and an N-terminal region comprising two α-helices connected by flexible linkers that extend into the downstream DNA channel and toward the RNAP active site, which is composed of β and β’ (Figure 9B). This interaction is consistent with the hypothesis that ML1026 is a part of actinobacterial RNAP.

ML1638 represents a distinct class of RNAP-associated proteins. It consists of an extended coiled-coil domain capped by a C-terminal zinc-ribbon motif and was strongly predicted to interact with β’ (size-corrected ipTM = 0.36), but not with β. The *M. smegmatis* homolog (MSMEG4306) has previously been structurally characterized^73^ and shown to be homologous to CT398 from *Chlamydia trachomatis* and HP0958 from *Helicobacter pylori*^73^. HP_0958 and CT398 are thought to interact with and stabilize RpoN^74,75^, a member of the activator-dependent σ^54^ family, which is absent in Actinobacteria^76^. Co-folding ML1638 with the *M. leprae* RNAP core enzyme (α₂ββ′ω) revealed a striking architecture in which the zinc-ribbon domain anchors the protein near β′ shelf and β′ jaw domains, partially overlapping the ML1026 binding site, while the elongated coiled-coil traverses the surface of RNAP across the swivel module (Figure 9C)^77,78^. The distal tip of the coiled-coil extends toward the canonical σ-factor binding region of RNAP. AlphaFold3 prediction of CT398 with *Chlamydia trachomatis* RNAP σ^N^-holoenzyme (α₂ββ′ωσ^N^) closely mirrors the mycobacterial RNAP-binding mode of ML1638. In the CT398 model, the distal coiled-coil tip contacts the core-binding domain (CBDa) of σ^54^ (Figure 9D), providing a structural rationale for previous proposals that CT398-family proteins regulate sigma-factor function. In Actinobacteria, where σ^54^ is absent, ML1638 may instead regulate the recruitment or positioning of other alternative sigma factors through direct interactions with the RNAP holoenzyme.

Collectively, our analysis revealed two previously unrecognized RNAP-binding factors. These two new factors, one of which is essential and broadly conserved in actinobacteria (ML1026), and one of which represents an evolutionarily widespread and understudied family of RNAP binding proteins (ML1638), expand our knowledge of mycobacterial RNAP and set the stage for future work in these and other bacterial genera.

### ML0986 may interact with DNA gyrase

DNA gyrase is an essential bacterial type II topoisomerase that introduces negative DNA supercoils required for both replication and transcription in bacteria^79^. DNA gyrase is the cellular target of (fluoro)quinolone antibiotics^80^ that trap the enzyme in its DNA-cleaved state, resulting in accumulation of double-stranded DNA breaks and ultimately bacterial cell death. Consequently, increased gyrase activity enhances quinolone-mediated DNA damage and accelerates killing. Our predicted PPIs identified ML0986, a 67-amino-acid DUF3046-family protein, as a predicted interaction partner of GyrA (size-corrected ipTM = 0.32), but not GyrB (size-corrected ipTM = 0.02), with a size-corrected ipTM similar to that of the GyrA-GyrB interaction (size-corrected ipTM = 0.33). Moreover, depletion of the ML0986 homolog in *M. tuberculosis* (Rv2738c) resulted in resistance to levofloxacin^12^, suggesting a functional and physical connection to gyrase. DUF3046 proteins are restricted to but widespread in Actinobacteria (∼50% of genomes), and are frequently encoded adjacent to RecA, a central player in DNA repair.

To understand how ML0986 may affect gyrase function, we co-folded the GyrA_2_GyrB_2_ heterotetramer with ML0986 (Figure 10A left panel). Gyrase is composed of a heterotetramer (GyrA_2_GyrB_2_) that forms three “gates” which undergo conformational changes that catalyze DNA strand passage (Figure 10A right panel). The DNA-gate, located at the GyrA–GyrB interface, binds and transiently cleaves one DNA duplex (the G-segment) to permit passage of a second duplex (the T-segment). ATP-dependent dimerization of GyrB forms the N-gate, which captures and directs the T-segment toward the DNA gate, while the C-gate (also referred to as the exit gate), formed by the GyrA dimer interface, opens to release the transported DNA duplex following strand passage. AlphaFold3 consistently positioned ML0986 at the C-gate (exit gate) of DNA gyrase (Figure 10A), where it occupies a pocket formed by the juxtaposed coiled-coil domains of the GyrA dimer and lies along the twofold symmetry axis of the complex (Figure 10B). Interestingly, the predicted binding site closely overlaps that of the CcdB gyrase toxin in *Escherichia coli* gyrase, despite ML0986 sharing no detectable sequence or structural similarity with CcdB. In the crystal structure of *E. coli* GyrA_2_-CcdB^81^, the CcdB homodimer binds the same GyrA coiled-coil region at the C-gate that is occupied by ML0986 in our model (Figure 10C). Because CcdB inhibits gyrase by preventing the conformational changes required for strand passage, this structural convergence suggests that ML0986 might similarly repress gyrase activity. However, the levofloxacin resistance of the *rv2738c* knockdown strain is consistent with ML0986/Rv2738c functioning as a gyrase activator, thus ML0986 binding in the C-gate may help modulate conformational transitions at the C-gate or regulate strand passage. Intriguingly, no mutations in *rv2738c* have been observed in clinical samples of *M. tuberculosis*, despite the substantial levofloxacin resistance associated with its depletion, suggesting it may play an important role in the presence of other stressors (e.g., DNA damage).

**Figure 10.**
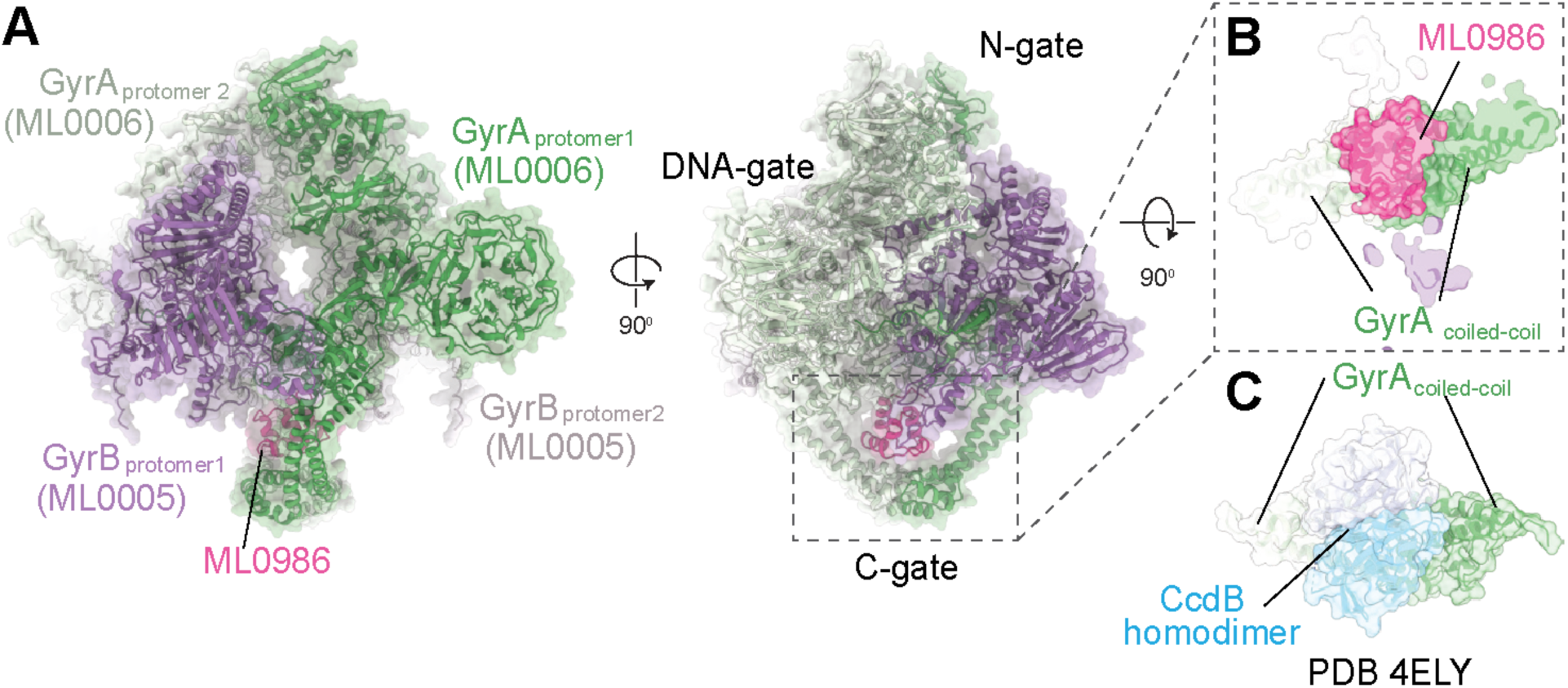
Predicted interaction of ML0986 with *M. leprae* DNA gyrase. **(A)** AlphaFold3 model of the *M. leprae* DNA gyrase heterotetramer. GyrA (ML0006) and GyrB (ML0005) protomers are shown as ribbon representations overlaid with transparent molecular surfaces and coloured individually. ML0986 (magenta) is positioned at the C-gate region of the enzyme. Orthogonal views reveal association of ML0986 with the coiled-coil domains of the GyrA dimer at the C-gate. **(B)** Close-up view of the ML0986-binding site following a 90° rotation relative to panel **A**. ML0986 (magenta) occupies a pocket formed by the juxtaposed GyrA coiled-coil domains (pale green and green) and is positioned along the twofold symmetry axis of the gyrase complex. **(C)** Structural comparison of the predicted ML0986-binding site with the crystal structure of the *Shigella flexneri*/*Aliivibrio fischeri* gyrase–CcdB complex (PDB: 4ELY^81^). The CcdB homodimer (pale cyan and cyan) binds the GyrA coiled-coil region (pale green and green) at a location analogous to that predicted for ML0986.

## PERSPECTIVE

In this manuscript, we show that combining the proteome-scale PPI network of the mycobacterial pathogen *M. leprae* predicted using pooled-AlphaFold3 with chemical genomics data from related organisms facilitates discovery of novel protein functions. We find that PPIs frequently underlie phenotypic correlations and demonstrate how associating a novel protein with well-characterized partners can accelerate and incentivize mechanistic dissection of its role. Although not all phenotypic correlations are driven by predicted PPIs, and not all proteins in predicted PPIs have correlated phenotypes, a significant fraction of proteins are predicted to have strong (28%) or intermediate (68%) strength interactions, and these interactions are broadly informative of gene function - gene pairs with high phenotypic correlations were 50- to 367- fold enriched for strong predicted PPIs.

The substantial overlap we observed between phenotypic correlations and predicted PPIs stands in contrast to observations in the eukaryotic yeast *Saccharomyces cerevisiae*, where deeply characterized PPI and genetic interaction (GI) networks were found to be largely distinct (i.e., <0.4% of GIs are due to PPIs)^82^. This discrepancy likely reflects fundamental differences in genome size, network density, and complex architecture between eukaryotes and bacteria^83^, but may also be due to differences in how PPI networks were determined. In contrast to the experimental approaches used to map yeast PPIs, our pooled-AlphaFold3 approach: 1) focuses on direct PPIs (i.e., two non-interacting proteins in the same complex would not be detected); 2) may have a lower false-positive rate; and 3) is heavily driven by co-evolutionary information, biasing our PPI predictions towards those that have functional implications. Ongoing studies using pooled-AlphaFold3 to predict the yeast PPI network (e.g., ref^84^) will establish the extent to which differences in the overlap between phenotypic (e.g., chemical genomics, or GIs) networks and PPI networks reflect fundamental differences between bacteria and eukaryotes.

In addition to specific interactions and gene functions, our predicted PPI data also revealed several broader paradigms. First, we find that GT-C family glycosyltransferases, which use lipid-linked sugar donors to build extracytoplasmic polysaccharides such as peptidoglycan and arabinogalactan, depend almost universally on PPIs to regulate their activity. In mycobacteria, our data and recent experimental studies find essential protein partners for numerous GT-Cs: Pmt (interacts with ML2687c, ML0810c, and ML1065), AftA (interacts with LpqZ), AftB (interacts with FecB and ML1720), AftC (interacts with ML1813c), AftD (interacts with TmaT), and potentially PimE (with ML0489). Second, we find that small proteins (which are often ignored despite their conservation across phyla) frequently play important and even essential roles in central processes. We identified ML1026 as an essential component of RNA polymerase, ML0986 as a DNA gyrase-interacting factor, ML1065 as a component of the protein mannosyltransferase complex, ML1347 as an essential co-factor of MtrP, ML0614 as an essential partner of MptB, and ML2141 as a partner of PgsA2, and confirmed several of these predictions with targeted experiments. Finally, and most generally, these data illustrate the limitations of traditional functional annotations and show how predicted PPIs can overcome them. For example, EphE, annotated as an epoxide hydrolase, appears to play a role in PIM synthesis and may use a radically different catalytic mechanism. The accuracy of our combined approach encourages the mechanistic dissection of such unusual gene functions.

Our choice of *M. leprae,* a reduced genome mycobacterium, as the target of our study was intentional. First, as a representative of the mycobacteria, it allowed us to demonstrate the utility of this approach in a relatively well-studied clade with major implications for human health. Second, its reduced genome minimized computational cost and the number of false-positive hits while retaining core mycobacterial processes. Finally, *M. leprae* is genetically intractable and can only be grown on nine-banded armadillos. By applying pooled-AlphaFold3 to this genome, we demonstrate the utility of our computational approach for understanding important but unculturable bacteria.

Our combined chemical-genomics and pooled-AlphaFold3 approach is broadly applicable. Tn-seq chemical-genomics data-sets are available for more than 60 bacteria genera^3^, and the increasing applicability of both CRISPRi and TnSeq to novel bacterial genomes (e.g., ref^85^) combined with the decreasing cost of DNA synthesis and next-generation sequencing suggests such datasets will continue to proliferate. Augmenting new and previously published chemical genomics screens with pooled-AlphaFold3 PPI networks will greatly accelerate mechanistic dissection of correlated phenotypes, driving studies of novel proteins and gene functions. In addition to chemical genomics, PPIs can also augment other genome-scale datasets, such as GIs^86,87^. Pooled-AlphaFold3 can also be used in the absence of experimental data, enabling initial studies of unculturable and/or intractable systems. Taken together, this manuscript establishes PPI prediction via pooled-AlphaFold3 as a major new tool for deciphering bacterial gene function and reveals numerous novel interactions that can be further pursued to better understand mycobacterial biology. To ensure wide access to this resource, all PPI data and predicted structures are publicly available through an interactive database (https://pebble.rockefeller.edu/pages/leprae_cofolding/).

## ACKNOWLEDGEMENTS

We thank Romas Kazlauskas for his advice on EphE and Michel Riviere for his insights on Pmt. We thank Stephanie Bueler and John Rubinstein for their assistance and advice for performing ORBIT and for sharing materials. We thank Joseph Adjei, Stefany Quinones Garcia, Abigail Garrett, Mariko Kanai, Benjamin Patty, Isabelle Seckler, Christina Stallings, and Junpei Xiao for running jobs using their personal AlphaFoldServer accounts. We thank Google and DeepMind for making AlphaFold3 freely available through <u>AlphaFoldServer.com</u>.

## METHODS

### Preparing pools

The genome of *M. leprae* TN was downloaded from Mycobrowser^88^ Release 5 (https://mycobrowser.epfl.ch/). Pools were designed as described in ref^5^. Briefly, proteins were pooled to minimize the number of pools required to co-fold all protein pairs at least once while keeping all jobs <5,000 tokens using a greedy algorithm available at https://github.com/horiatodor/pooled-af3. We excluded ML1191 (*fas*), because its large size (3,076aa) precluded screening against all other proteins.

### Running AlphaFold3

All 17,861 comprehensive pools were successfully run using default settings on the AlphaFold3 server (https://alphafoldserver.com/). The descriptions for all of the pools are available in Table S1.

### Analysis of AlphaFold3 runs

All AlphaFold3 runs were downloaded. Raw ipTM scores were extracted from the “summary_confidences” files for all 5 models and averaged. The ipTM of protein pairs that appeared in multiple pools was averaged across all pools. Analysis code is available at https://github.com/horiatodor/pooled-af3.

### Size correction of ipTM scores

ipTM size correction was performed as previously described^5^. To determine the effect of summed protein size on ipTM, we performed a robust linear regression using the robustbase::lmrob function, which computes a MM-type regression estimator (a robust and efficient estimator with a ∼50% breakdown point and 95% efficiency). A robust estimator was used to mitigate the influence of true-positive interactions on the regression. The linear regression was performed on the square root of the summed size of the two proteins (in amino acids) to calculate an “expected ipTM” for each protein pair. For the *M. leprae* dataset, expected_ipTM = -0.035824446 + 0.004441653*sqrt(aa_in_protein1 + aa_in_protein2), values consistent with our previous dataset^5^. Expected ipTM was subtracted from the observed ipTM to generate the “size-corrected ipTM”. Code for implementing this correction is available at https://github.com/horiatodor/pooled-af3.

### STRING data

STRING data related to taxid: 272631 was downloaded on April 24th, 2025 from https://string-db.org/. Only the experimental channel was used for benchmarking PPIs.

#### Data Retrieval and Processing

*M. smegmatis* CRISPRi. Data and correlations were taken directly from ref^10^.
*M. tuberculosis* CRISPRi. Raw data from ref^12^ was retrieved from https://github.com/rock-lab/CGI_nature_micro_2022. Data was processed, corrected, and correlations were quantified as previously described^89^.
*C. glutamicum* Tn-Seq. Processed per-gene fitness data was retrieved from the Fig. 2—source data 1 file of ref^11^. Data was processed, corrected, and correlations were quantified as previously described^89^.

### M. smegmatis ephE knockout mutant

*M. smegmatis ephE* deletion was generated using Ts-SacB methodology^90^. All strains were grown in 7H9-OADC broth (BD, Difco) supplemented with 0.05% Tween 80 at 37°C; where required, kanamycin (25 µg/mL) and hygromycin (50 µg/mL) were added to the culture medium. Briefly, 717-bp of the coding sequence of *ephE,* bracketed between a *Pst*I and an *Age*I restriction site, was deleted and replaced by a kanamycin resistance cassette. Gene replacement was confirmed by PCR using primers located outside the allelic exchange substrate (Figure S2D). The *ephE* deletion mutant was complemented with a wild-type copy of the *ephE* gene from *M. tuberculosis* (*Rv3670*) expressed under control of the *hsp60* promoter from the replicative plasmid pVV16^91^ or under control of its own promoter from the integrative plasmid pNIP40b^92^.

### Whole cell radiolabeling experiments and lipid analyses

Radiolabeling of whole *M. smegmatis* cultures grown to an OD600 ∼ 0.4 with [1,2-^14^C]acetic acid (0.5 mCi/mL; specific activity, 57 Ci/mol, NEN Radiochemicals) was performed in 7H9-OADC-Tween 80 medium for 24 hours at 37°C with shaking. Total lipids from bacterial cells were extracted with CHCl_3_/CH_3_OH (1:2, vol:vol) for one night followed by two overnight extractions with CHCl_3_/CH_3_OH (2:1, vol:vol). [1,2-^14^C] acetic acid-derived lipids were analyzed by thin-layer chromatography (TLC) on aluminum-backed silica gel 60-precoated plates F_254_ (E. Merck) and radiolabeled products were semi-quantified using a Sapphire Biomolecular Imager.

### Generation of endogenously tagged Pmt

*M. smegmatis Pmt* strains were generated using ORBIT^93^. All strains were grown in Middlebrook 7H9 media (Difco) (supplemented with 0.5% w/v BSA, 11 mM dextrose, 14 mM NaCl, 0.2 % w/v glycerol, 0.05% w/v Tween-80), and appropriate antibiotics. An initial strain (*M. smegmatis* mc^2^155) containing pKM461 (Addgene #108320), was used to inoculate a 10 mL starter culture which was grown to saturation at 37°C. This culture was used to inoculate a second 25 mL culture, which was grown to an OD of 0.5. 500 ng/mL of anhydrotetracycline (SupelCo #37919) was added to the culture to allow expression of the Che9c phage annealase and the Bxb1 integrase. pKM461 was a gift from Kenan Murphy. After growing for another 3.5 hours, the culture was cooled on ice for 10 minutes and 20 mL were harvested by centrifugation of 3000x*g*. The cells were then washed twice in sterile, ice cold 10% glycerol. To tag endogenous Pmt (MSMEG_5447), 380 μL of electrocompetent cells were mixed with 200 ng of the payload plasmid pSAB41 and 1 μg of the targeting DNA oligomer (IDT DNA Ultramer) (5’-CCCGATCCTGACGGGCCTGCCGATCTCACAGACCACCTGGAACCTGCAGATCTGGTTGCCGAGCTGGCGCGGTTTGTCTGGTCAACCACCGCGGTCTCAGTGGTGTACGGTACAAACCTAGCTTCTGTTTGTTCCGGCGTCCTCCCCTATCAGTAGTTCTGTATTGGTTCCCGTCCTCGAAACGGACA -3’). The cells were then electroporated in an ice-cold 0.2 cm gap cuvette (BioRad #1652086) and allowed to recover in 2 mL antibiotic free Middlebrook 7H9 media overnight. 200 μL of the culture was then plated on a 7H10 plate containing 50 μg/mL hygromycin B (Goldbio #H-270-1) and incubated at 37°C for 4 to 5 days. Positive colonies were verified for the chromosomal insertion using PCR or whole genome sequencing (Plasmidsaurus) and stored at -80°C as glycerol stocks.

### Expression and purification of endogenously tagged Pmt

A Pmt-3xFLAG glycerol stock was streaked on an LB agar plate containing 50 μg/mL hygromycin B and incubated at 37°C for 2 days. The WT strain, containing only pKM461, was streaked on LB agar containing 25 μg/mL kanamycin. Colonies were picked and used to inoculate a 20 mL starter culture which was grown at 37°C at 180 RPM for two days. This culture was used to inoculate 6 L of Middlebrook 7H9 media and which were grown at 37°C shaking at 180 RPM for two days. The cells were then harvested with centrifugation at 6000x*g* for 10 minutes in an Avanti J-E centrifuge (Beckman Coulter) and flash frozen and stored at - 80°C until further use. After thawing, the cells were resuspended in Lysis Buffer (50 mM Tris pH 7.5, 100 mM NaCl, 20 mM MgSO_4,_ and 5 mM TCEP) using 5 mL buffer per gram of cells. The solution was then supplemented with 20 mg/mL DNase1, 0.2 μg/mL phenylmethylsulfonyl fluoride (PMSF), and EDTA-free cOmplete protease inhibitor cocktail (1 tablet per liter) (Roche) and lysed with 4 passages through an Emulsiflex C3 homogenizer (Avastin) operating at a pressure of 10,000 – 15,000 PSI. Cell debris and unlysed cells were pelleted by centrifugation at 3000x*g* in a Centrifuge 5510 R (Eppendorf) for 10 minutes. Membranes were then pelleted by centrifugation at 185,600x*g* in a Type 45 Ti Rotor (Beckman Coulter) for 1 hour at 4°C. The membrane pellets were resuspended in Lysis Buffer (15 mL per gram of membrane) and flash frozen until further use. After thawing, n-docyl-β-D-maltopyranoside (DDM) (Inalco #1758-1350) was added to the solution to a final concentration of 1% and stirred at 4°C for 3 hours. The solution was then clarified with centrifugation at 185,600x*g* for 30 minutes at 4°C. The supernatant was then applied to a gravity column packed with 200 μL of α-FLAG G1 Affinity resin (GenScript #L00432). The column was washed with 10 CV of Wash Buffer 1 (50 mM Tris pH 7.5, 150 mM NaCl, and 0.02% DDM) and 10 CV of Wash Buffer 2 (50 mM Tris pH 7.5, 300 mM NaCl, and 0.02% DDM). Protein was eluted with 4 CV of Wash Buffer 1 containing 150 μg/mL of 3xFLAG peptide (GLPBIO #GP10149). The elution fraction was then concentrated ∼10-fold using an Amicon Ultra Centrifugal Filter (Millipore Sigma #UFC501024) and 10 to 15 μg of protein (30 - 40 μL) was mixed with 6x reducing Laemmli SDS Sample Buffer (Boston BioProducts #BP-111R) and analyzed with SDS-PAGE using 4-20% Mini-PROTEAN TGX Precast Protein Gels (BioRad #456109), followed by staining with Instantblue Coomassie Protein Stain (Adcam #ab119211).

### Mass spectrometry protein identification

After SDS-PAGE analysis, bands of interest were excised and transferred to 20 μL of sterile 1% acetic acid and stored on dry ice. Mass spectrometry analysis was performed by The Protein Facility of the Iowa State University Office of Biotechnology. The samples were thawed and reduced with DTT, the cysteine residues were modified with iodoacetamide, and digested overnight with trypsin/Lys-C. The digestion was stopped with the addition of formic acid, and the samples were dried in a SpeedVac. Samples were then desalted using a BioPureSPN MiniPROTO 300 C18 column (Nest Group Bio) before drying again in a SpeedVac. PRTC standard (Pierce #88320) was spiked into the sample to serve as an internal control. The peptides were then separated by liquid chromatography on an EASY-Spray PepMap Neo 75 μm/150 μm x 150 mm column (ThermoScientific #ES75150PN), operated on a Vanquish NEO HPLC system coupled with an Easy-Spray Source (ThermoScientific). The column was washed with Buffer A (0.1% formic acid) and peptides were eluted with Buffer B (80% acetonitrile, 0.1% formic acid). The peptides were analyzed by MS/MS on an Orbitrap Astral (ThermoScientific) by fragmenting each peptide. The resulting intact and fragmentation pattern was compared to a theoretical fragmentation pattern to find usable peptides for protein identification. The PRTC areas were used to normalize data between samples. The resulting peptides were searched using CHIMERYS^94^ against the *M. smegmatis* mc^2^155 genome and analyzed with ProteomeDiscover (ThermoScientific).

## DATA AVAILABILITY

All data derived from alphafoldserver.com is available through an interactive database (https://pebble.rockefeller.edu/pages/leprae_cofolding/), subject to AlphaFold Server terms of service, which can be found at https://alphafoldserver.com/terms.

## SUPPLEMENTARY FIGURES

**Figure S1.**
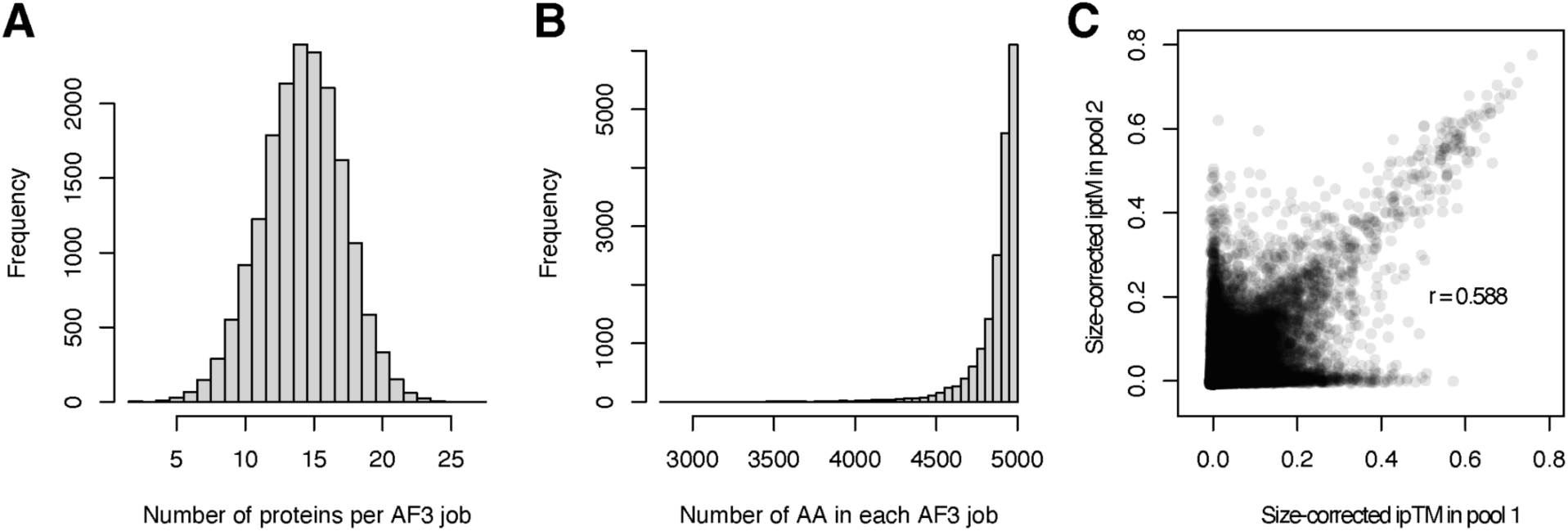
**(A)** Histogram of number of proteins in each job (Table S1). **(B)** Histogram of the total size of each of 17,861 pools (Table S1). **(C)** Pooled-AlphaFold3 ipTM scores are reproducible. Size-corrected ipTM scores for the 533,898 protein pairs that appear in multiple pools are similar (Pearson’s r = 0.588, 533,898 protein pairs).

**Figure S2.**
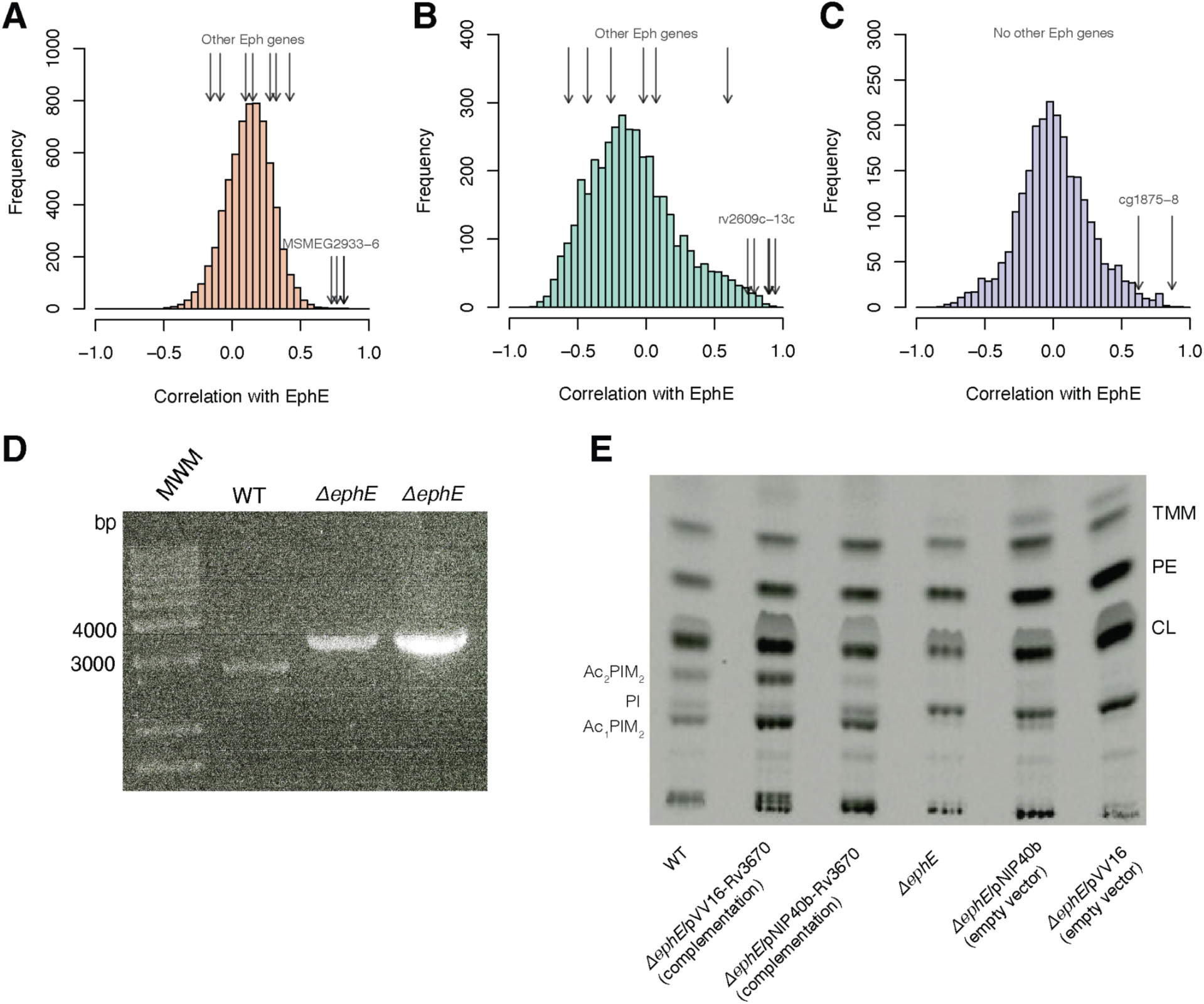
Additional data regarding EphE. (A-C) Histogram of phenotypic correlations between *ephE* (ML2297/MSMEG6184/Rv3670/Cg0358) and all other genes in (A) *M. smegmatis*; (B) *M. tuberculosis*; and (C) *C. glutamicum* showing strong phenotypic correlations between knockouts/knockdowns of *ephE* and those of genes in the *pimA* operon (ML0451-5c/MSMEG2933-6/Rv2609-13c/Cg1875-8), but not to other epoxide hydrolases. EphE and all other proteins in the *M. leprae* data. Opacity of points represents the phenotypic correlations in *M. smegmatis* (orange), *M. tuberculosis* (green), and *C. glutamicum* (purple). **(D)** Gene replacement at the *ephE* locus of *M. smegmatis*. Allelic replacement at the *ephE* locus was confirmed by PCR using primers located outside the allelic exchange substrate. The expected sizes of the PCR products are 3,000-bp in the WT strain and 3,500-bp in the *ΔephE* deletion mutant. Two individual mutant clones are shown. **(E)** Total [1,2-^14^C]acetate-derived lipids prepared from WT, *ΔephE, ΔephE* carrying an empty pVV16 plasmid or an empty pNIP40 plasmid (“empty vector”), and *ΔephE* complemented mutant expressing *ephE* from *M. tuberculosis* from pVV16 or pNIP40b (“complementation”) were analyzed by TLC using CHCl_3_/CH_3_OH/H_2_O (65:25:4, by vol) as the eluent. The same total counts (5,000 dpm) of each lipid sample were loaded per lane. TMM, trehalose monomycolates; PE, phosphatidylethanolamine; CL, cardiolipin; PI, phosphatidy-myo-inositol; Ac₁PIM₂, triacylated phosphatidylinositol dimannosides; Ac₂PIM₂, tetraacylated phosphatidylinositol dimannosides.

**Figure S3.**
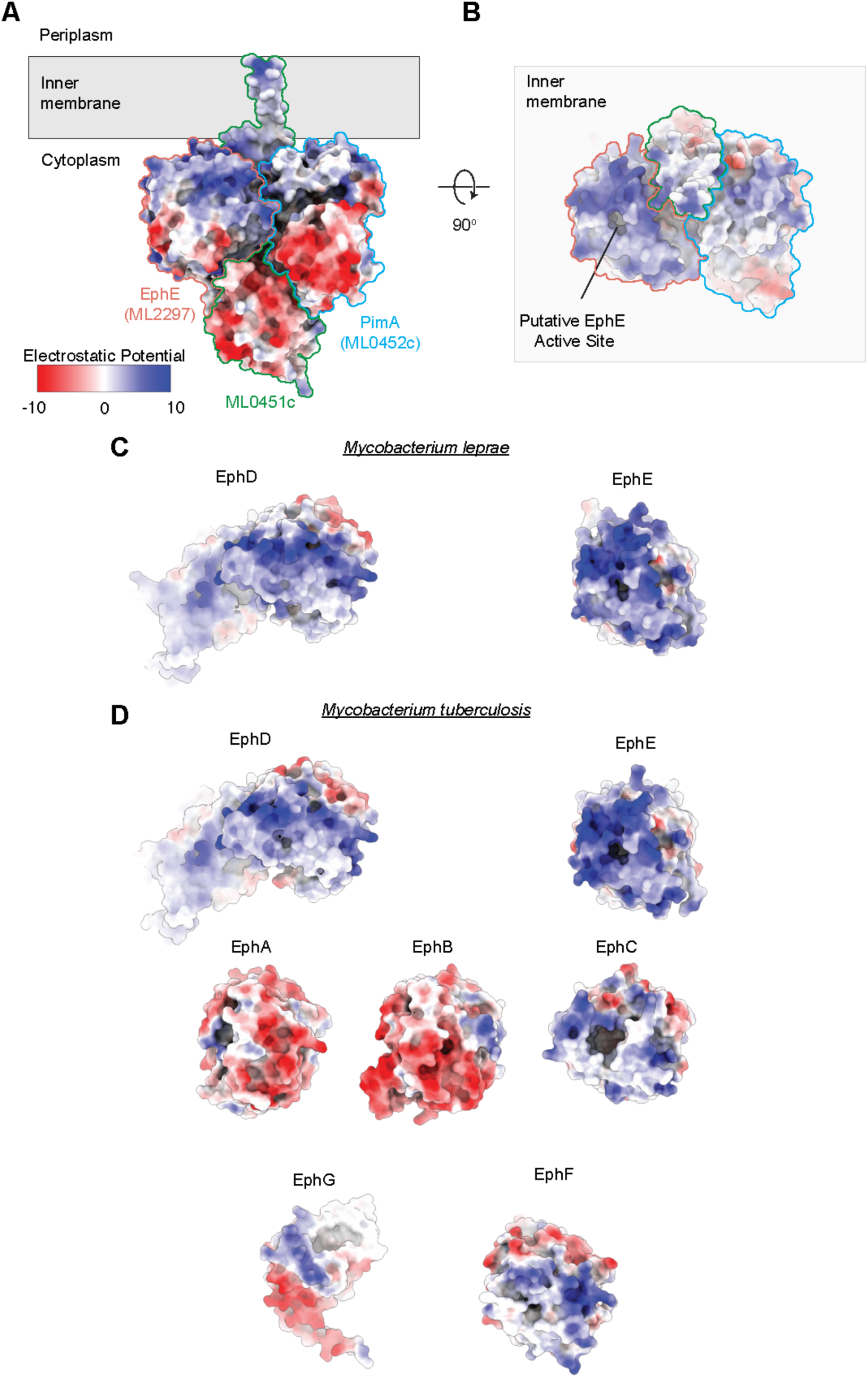
Electrostatic surfaces of the PimA–ML0451c–EphE complex and comparison with the mycobacterial epoxide hydrolase family. **(A)** Electrostatic surface representation of the AlphaFold3-predicted *M. leprae* PimA–ML0451c–EphE complex viewed parallel to the inner membrane. Electrostatic potential is mapped onto the molecular surface, with red and blue denoting regions of negative and positive electrostatic potential, respectively. The boundaries of PimA (ML0452c, blue), ML0451c (green), and EphE (ML2297, salmon) are outlined. **(B)** Cytoplasmic view of the complex following a 90° rotation relative to (A). The putative EphE active site is located within a positively charged surface patch oriented toward the inner membrane. **(C)** Electrostatic surface representations of *M. leprae* EphD and EphE. Whereas EphD exhibits a mixed distribution of positive and negative surface potentials surrounding the putative active site, EphE is characterized by a pronounced electropositive surface pocket. **(D)** Electrostatic surface comparison of the seven annotated epoxide hydrolases from *Mycobacterium tuberculosis*. EphE displays a strongly electropositive active-site environment relative to EphA, EphB, EphC, EphD, EphF and EphG.

**Figure S4.**
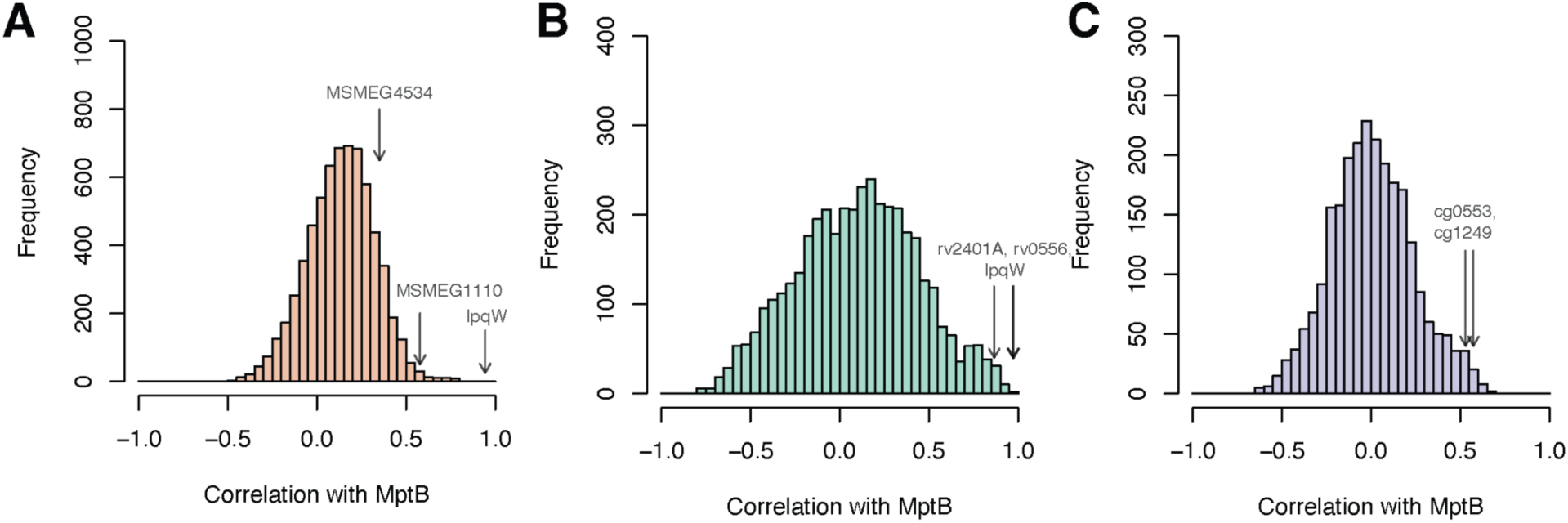
Additional data regarding MptB. (A-C) Histogram of phenotypic correlations between *mptB* and all other genes in (A) *M. smegmatis*; (B) *M. tuberculosis*, and; (C) *C. glutamicum* showing strong phenotypic correlations between knockouts/knockdowns of *mptB* and those of *lpqW,* and *ML0614* and *ML2271* homologs.

**Figure S5.**
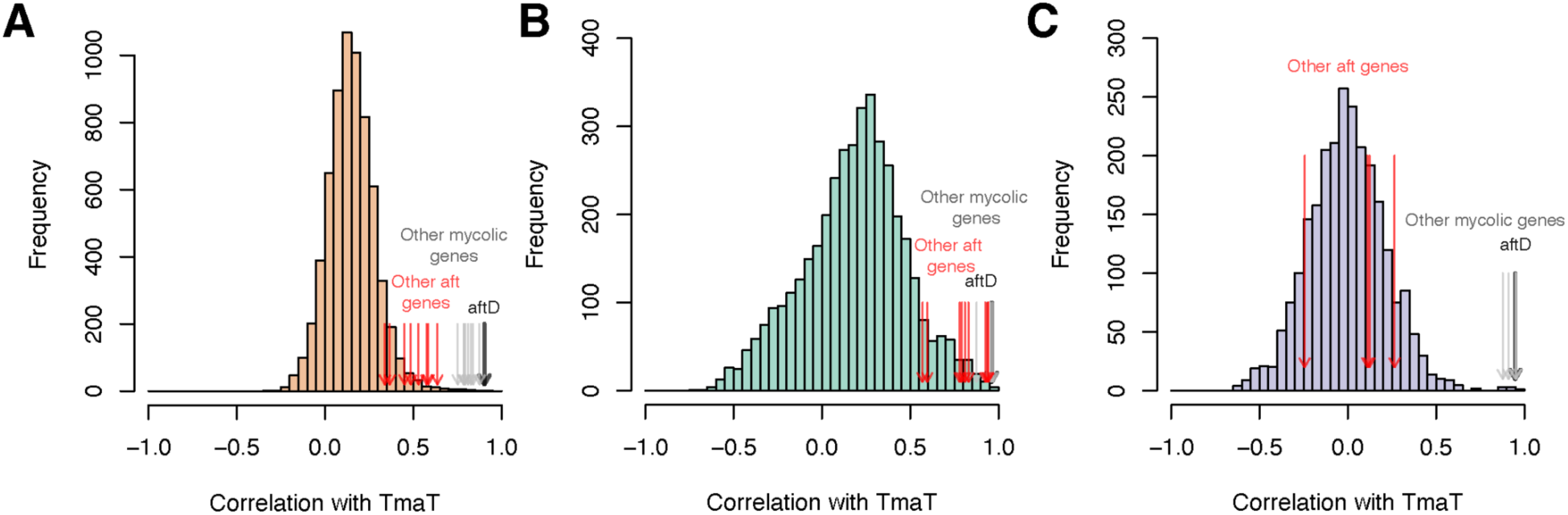
Additional data regarding TmaT. (A-C) Histogram of phenotypic correlations between *tmaT* and all other genes in (A) *M. smegmatis*; (B) *M. tuberculosis*, and; (C) *C. glutamicum* showing strong phenotypic correlation to *aftD* (dark black arrow) and other mycolic acid synthesis genes (*MSMEG6143, mmpA, accD5, MSMEG0834, mmpL3, pks13, MSMEG0311,* and *accD4*, grey arrows) but not to other arabinogalactan synthesis genes (*aftA, dprE2, wbbL1, glfT2, glfT1, aftB, aftC,* and *aftH*, red arrows).

**Figure S6.**
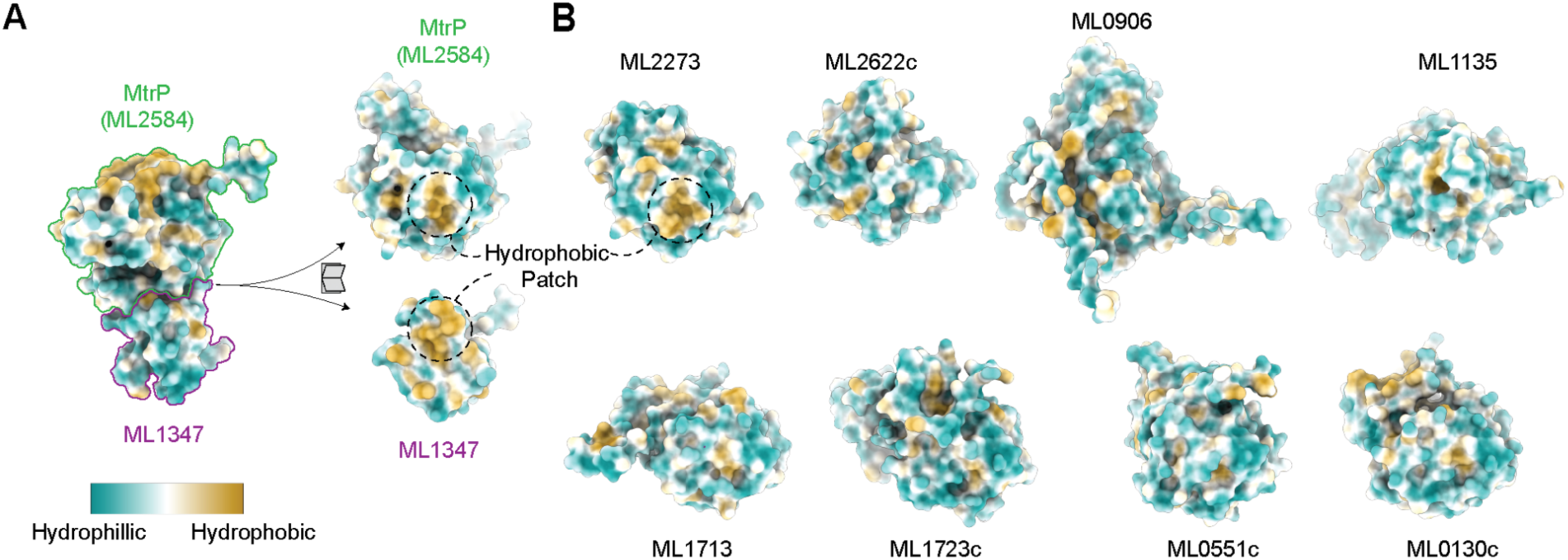
Conserved hydrophobic interaction surface in MtrP-associated proteins. **(A)** Surface hydrophobicity mapping of the AlphaFold3-predicted MtrP–ML1347 complex from *M. leprae*. Hydrophilic and hydrophobic regions are colored cyan and gold, respectively. MtrP (ML2584) and ML1347 exhibit complementary hydrophobic surface patches at the predicted heterodimer interface. The hydrophobic patch on ML1347 is highlighted. **(B)** Surface hydrophobicity representations of the *M. leprae* methyltransferases MenG (ML2273), ML2622c, ML0906, ML1135, ML1713, ML1723c, ML0551c and ML0130c. Among these proteins, ML2273 is predicted to interact with ML1347 and possesses a prominent hydrophobic surface patch analogous to that observed at the MtrP–ML1347 interface.

**Figure S7.**
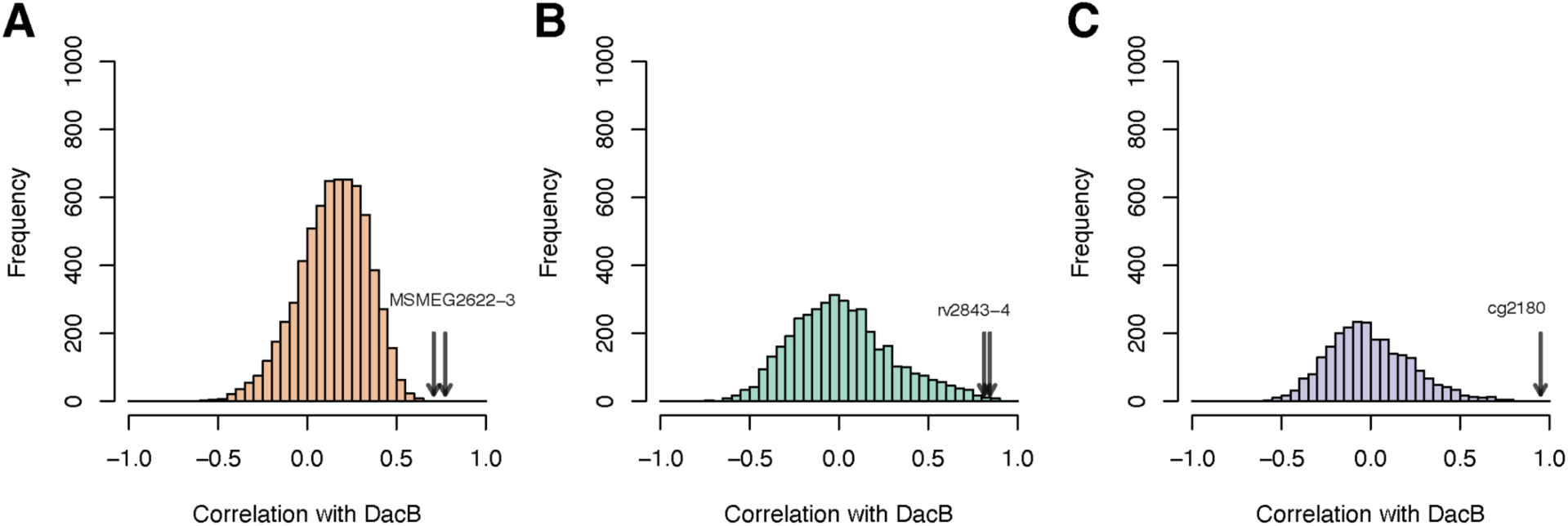
Additional data regarding DacB. Histogram of phenotypic correlations between *dacB* and all other genes in (A) *M. smegmatis*; (B) *M. tuberculosis*, and; (C) *C. glutamicum* showing strong phenotypic correlation to *ML1560-1* homologs.

**Figure S8.**
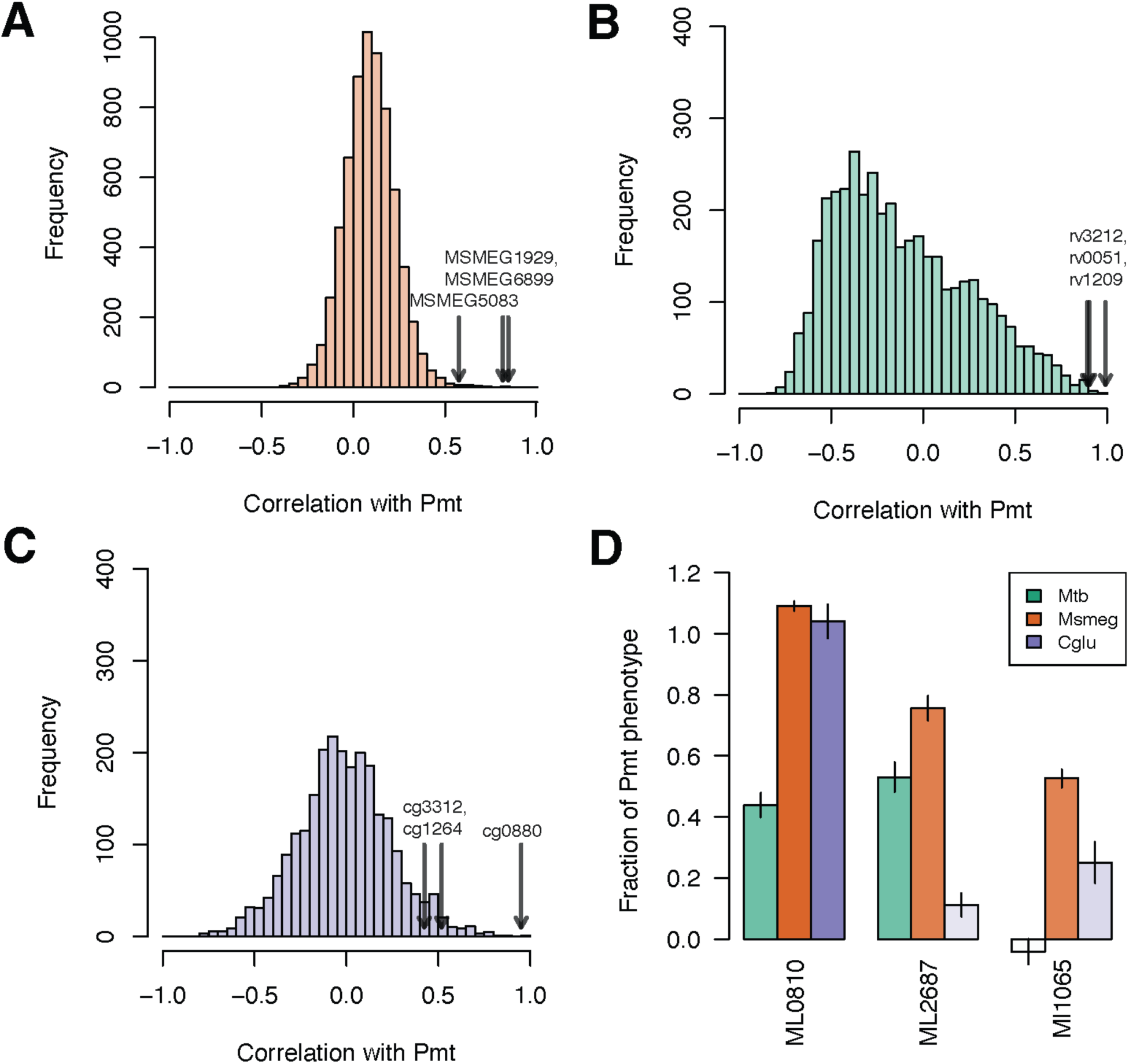
Additional data regarding Pmt. (A-C) Histogram of phenotypic correlations between *pmt* and all other genes in (A) *M. smegmatis*; (B) *M. tuberculosis*, and; (C) *C. glutamicum* showing strong phenotypic correlation to *ML0810* and *ML2687c* homologs, and a weaker correlation to *ML1065* homologs (dark black arrows). **(D)** Relative phenotypes of Pmt-complex components in all three bacteria show a clear order - ML0810c is the most important, followed closely by ML2687c, and ML1065 seems less important.

## SUPPLEMENTARY TABLES

Table S1 - Description of pools.

Table S2 - Matrix of size corrected ipTMs for all protein pairs (1609×1609).

Table S3 - Homologs of *M. leprae* proteins in *M. tuberculosis, M. smegmatis,* and *C. glutamicum*.

Table S4 - Enrichment of strong, intermediate, and weak predicted PPIs in phenotypically correlated gene pairs in *M. tuberculosis, M. smegmatis,* and *C. glutamicum* chemical genomics data.

Table S5 - Mass spectrometry data for Pmt (MSMEG5447) pulldown.

## REFERENCES

1. Vanni, C., Schechter, M.S., Acinas, S.G., Barberán, A., Buttigieg, P.L., Casamayor, E.O., Delmont, T.O., Duarte, C.M., Eren, A.M., Finn, R.D., et al. (2022). Unifying the known and unknown microbial coding sequence space. Elife 11, e67667.

2. Nichols, R.J., Sen, S., Choo, Y.J., Beltrao, P., Zietek, M., Chaba, R., Lee, S., Kazmierczak, K.M., Lee, K.J., Wong, A., et al. (2011). Phenotypic landscape of a bacterial cell. Cell 144, 143–156.

3. Price, M.N., Wetmore, K.M., Jordan Waters, R., Callaghan, M., Ray, J., Liu, H., Kuehl, J.V., Melnyk, R.A., Lamson, J.S., Suh, Y., et al. (2018). Mutant phenotypes for thousands of bacterial genes of unknown function. Preprint, 10.1038/s41586-018-0124-0 https://doi.org/10.1038/s41586-018-0124-0.

4. Todor, H., Silvis, M.R., Osadnik, H., and Gross, C.A. (2021). Bacterial CRISPR screens for gene function. Curr. Opin. Microbiol. 59, 102–109.

5. Todor, H., Kim, L.M., Jänes, J., Burkhart, H.N., Darst, S.A., Beltrao, P., and Gross, C.A. (2026). Predicting the protein interaction landscape of a free-living bacterium with pooled-AlphaFold3. Mol. Syst. Biol. 10.1038/s44320-026-00189-7.

6. DeJesus, M.A., Gerrick, E.R., Xu, W., Park, S.W., Long, J.E., Boutte, C.C., Rubin, E.J., Schnappinger, D., Ehrt, S., Fortune, S.M., et al. (2017). Comprehensive essentiality analysis of the Mycobacterium tuberculosis genome via saturating transposon Mutagenesis. MBio 8, e02133–16.

7. Marmiesse, M., Brodin, P., Buchrieser, C., Gutierrez, C., Simoes, N., Vincent, V., Glaser, P., Cole, S.T., and Brosch, R. (2004). Macro-array and bioinformatic analyses reveal mycobacterial “core” genes, variation in the ESAT-6 gene family and new phylogenetic markers for the Mycobacterium tuberculosis complex. Microbiology 150, 483–496.

8. Dulberger, C.L., Rubin, E.J., and Boutte, C.C. (2020). The mycobacterial cell envelope - a moving target. Nat. Rev. Microbiol. 18, 47–59.

9. Cole, S.T., Eiglmeier, K., Parkhill, J., James, K.D., Thomson, N.R., Wheeler, P.R., Honoré, N., Garnier, T., Churcher, C., Harris, D., et al. (2001). Massive gene decay in the leprosy bacillus. Nature 409, 1007–1011.

10. Herrera, N., Todor, H., Kim, L.M., Burkhart, H.N., Billings, E., Fay, A., Warner, T.C., Lee, S.Y., Sayegh, N.Y., Bosch, B., et al. (2025). The phenotypic landscape of the mycobacterial cell. Microbiology.

11. Sher, J.W., Lim, H.C., and Bernhardt, T.G. (2020). Global phenotypic profiling identifies a conserved actinobacterial cofactor for a bifunctional PBP-type cell wall synthase. Elife 9. 10.7554/eLife.54761.

12. Li, S., Poulton, N.C., Chang, J.S., Azadian, Z.A., DeJesus, M.A., Ruecker, N., Zimmerman, M.D., Eckartt, K.A., Bosch, B., Engelhart, C.A., et al. (2022). CRISPRi chemical genetics and comparative genomics identify genes mediating drug potency in Mycobacterium tuberculosis. Nat Microbiol 7, 766–779.

13. Neupane, P., Liu, J., and Cheng, J. (2026). Improving AlphaFold3 by engineering MSA and template inputs. bioRxivorg, 2026.04.22.720119. 10.64898/2026.04.22.720119.

14. Vissa, V.D., and Brennan, P.J. (2001). The genome of Mycobacterium leprae: a minimal mycobacterial gene set. Genome Biol. 2, REVIEWS1023.

15. Bansal-Mutalik, R., and Nikaido, H. (2014). Mycobacterial outer membrane is a lipid bilayer and the inner membrane is unusually rich in diacyl phosphatidylinositol dimannosides. Proc. Natl. Acad. Sci. U. S. A. 111, 4958–4963.

16. Šarkan, M., Forbak, M., Brown, C.M., Gilleron, M., Angala, S.K., De, K., Tymčuková, V., Gašparovič, H., Guerin, M.E., Nigou, J., et al. (2026). Essential role of MptB in the biosynthesis of phosphatidylinositol mannosides, lipomannan and lipoarabinomannan in mycobacteria. J. Biol. Chem. 302, 113077.

17. Global tuberculosis report 2024 (2024). https://www.who.int/publications/i/item/9789240101531.

18. Venkatesan, K., Rual, J.-F., Vazquez, A., Stelzl, U., Lemmens, I., Hirozane-Kishikawa, T., Hao, T., Zenkner, M., Xin, X., Goh, K.-I., et al. (2009). An empirical framework for binary interactome mapping. Nat. Methods 6, 83–90.

19. Szklarczyk, D., Kirsch, R., Koutrouli, M., Nastou, K., Mehryary, F., Hachilif, R., Gable, A.L., Fang, T., Doncheva, N.T., Pyysalo, S., et al. (2023). The STRING database in 2023: protein-protein association networks and functional enrichment analyses for any sequenced genome of interest. Nucleic Acids Res. 51, D638–D646.

20. Muro, E.M., Mah, N., Moreno-Hagelsieb, G., and Andrade-Navarro, M.A. (2011). The pseudogenes of Mycobacterium leprae reveal the functional relevance of gene order within operons. Nucleic Acids Res. 39, 1732–1738.

21. Lissner, R., Franklin, A., Benedict, S.T., Charitou, V., Speer, A., Knol, J., Jimenez, C., Moynihan, P.J., Nejentsev, S., Kuijl, C., et al. (2026). A new role for lipoproteins LpqZ and FecB in orchestrating mycobacterial cell envelope biogenesis. MBio 17, e0211925.

22. Klevorn, T., Brown, C., Hardy, C.D., Cuthbert, B.J., Spencer, A., Jinich, A., Chan, L., Angala, S.K., Manzer, J., Mendoza, J., et al. (2026). A periplasmic protein complex mediates arabinofuranosyltransferase activity and intrinsic drug resistance in Mycobacterium tuberculosis. Sci. Adv. 12, eaec5100.

23. Chen, J., Fruhauf, A., Fan, C., Ponce, J., Ueberheide, B., Bhabha, G., and Ekiert, D.C. (2023). Structure of an endogenous mycobacterial MCE lipid transporter. Nature 620, 445–452.

24. Boutte, C.C., Baer, C.E., Papavinasasundaram, K., Liu, W., Chase, M.R., Meniche, X., Fortune, S.M., Sassetti, C.M., Ioerger, T.R., and Rubin, E.J. (2016). A cytoplasmic peptidoglycan amidase homologue controls mycobacterial cell wall synthesis. Elife 5, e14590.

25. Wu, K.J., Boutte, C.C., Ioerger, T.R., and Rubin, E.J. (2019). Mycobacterium smegmatis HtrA blocks the toxic activity of a putative cell wall amidase. Cell Rep. 27, 2468–2479.e3.

26. Rahlwes, K.C., Osman, S.H., and Morita, Y.S. (2020). Role of LmeA, a Mycobacterial periplasmic protein, in maintaining the mannosyltransferase MptA and its product lipomannan under stress. mSphere 5. 10.1128/mSphere.01039-20.

27. Rainczuk, A.K., Yamaryo-Botte, Y., Brammananth, R., Stinear, T.P., Seemann, T., Coppel, R.L., McConville, M.J., and Crellin, P.K. (2012). The lipoprotein LpqW is essential for the mannosylation of periplasmic glycolipids in Corynebacteria. J. Biol. Chem. 287, 42726–42738.

28. Funck, T., McGowen, K., Sullivan, M.R., Zinga, S., Wolf, I.D., Nurjadi, D., Denkinger, C.M., and Rubin, E.J. (2026). Ribosomal protection as a linezolid resistance mechanism in Mycobacterium abscessus. Antimicrob. Agents Chemother, e0160525.

29. Jinich, A., Zaveri, A., DeJesus, M.A., Spencer, A., Almada-Monter, R., Flores-Bautista, E., Smith, C.M., Sassetti, C.M., Rock, J.M., Ehrt, S., et al. (2025). The Mycobacterium tuberculosis transposon sequencing database (MtbTnDB): A large-scale guide to genetic conditional essentiality. Mol. Microbiol. 124, 91–101.

30. Rock, J.M., Hopkins, F.F., Chavez, A., Diallo, M., Chase, M.R., Gerrick, E.R., Pritchard, J.R., Church, G.M., Rubin, E.J., Sassetti, C.M., et al. (2017). Programmable transcriptional repression in mycobacteria using an orthogonal CRISPR interference platform. Nat Microbiol 2, 16274.

31. Oh, E., Becker, A.H., Sandikci, A., Huber, D., Chaba, R., Gloge, F., Nichols, R.J., Typas, A., Gross, C.A., Kramer, G., et al. (2011). Selective ribosome profiling reveals the cotranslational chaperone action of trigger factor in vivo. Cell 147, 1295–1308.

32. Clarke, O.B., Tomasek, D., Jorge, C.D., Dufrisne, M.B., Kim, M., Banerjee, S., Rajashankar, K.R., Shapiro, L., Hendrickson, W.A., Santos, H., et al. (2015). Structural basis for phosphatidylinositol-phosphate biosynthesis. Nat. Commun. 6, 8505.

33. Morii, H., Ogawa, M., Fukuda, K., Taniguchi, H., and Koga, Y. (2010). A revised biosynthetic pathway for phosphatidylinositol in Mycobacteria. J. Biochem. 148, 593–602.

34. Korduláková, J., Gilleron, M., Mikusova, K., Puzo, G., Brennan, P.J., Gicquel, B., and Jackson, M. (2002). Definition of the first mannosylation step in phosphatidylinositol mannoside synthesis. PimA is essential for growth of mycobacteria: PimA IS ESSENTIAL FOR GROWTH OF MYCOBACTERIA. J. Biol. Chem. 277, 31335–31344.

35. Sancho-Vaello, E., Albesa-Jové, D., Rodrigo-Unzueta, A., and Guerin, M.E. (2017). Structural basis of phosphatidyl-myo-inositol mannosides biosynthesis in mycobacteria. Biochim. Biophys. Acta Mol. Cell Biol. Lipids 1862, 1355–1367.

36. Murat, D., Quinlan, A., Vali, H., and Komeili, A. (2010). Comprehensive genetic dissection of the magnetosome gene island reveals the step-wise assembly of a prokaryotic organelle. Proc. Natl. Acad. Sci. U. S. A. 107, 5593–5598.

37. Giganti, D., Albesa-Jové, D., Urresti, S., Rodrigo-Unzueta, A., Martínez, M.A., Comino, N., Barilone, N., Bellinzoni, M., Chenal, A., Guerin, M.E., et al. (2015). Secondary structure reshuffling modulates glycosyltransferase function at the membrane. Nat. Chem. Biol. 11, 16–18.

38. Ozhelvaci, F., and Steczkiewicz, K. (2025). Α/β hydrolases: Toward unraveling entangled classification. Proteins 93, 855–870.

39. Bosch, B., DeJesus, M.A., Poulton, N.C., Zhang, W., Engelhart, C.A., Zaveri, A., Lavalette, S., Ruecker, N., Trujillo, C., Wallach, J.B., et al. (2021). Genome-wide gene expression tuning reveals diverse vulnerabilities of M. tuberculosis. Cell 184, 4579–4592.e24.

40. Madacki, J., Kopál, M., Jackson, M., and Korduláková, J. (2021). Mycobacterial epoxide hydrolase EphD is inhibited by urea and thiourea derivatives. Int. J. Mol. Sci. 22, 2884.

41. Crellin, P.K., Kovacevic, S., Martin, K.L., Brammananth, R., Morita, Y.S., Billman-Jacobe, H., McConville, M.J., and Coppel, R.L. (2008). Mutations in pimE restore lipoarabinomannan synthesis and growth in a Mycobacterium smegmatis lpqW mutant. J. Bacteriol. 190, 3690–3699.

42. Kovacevic, S., Anderson, D., Morita, Y.S., Patterson, J., Haites, R., McMillan, B.N.I., Coppel, R., McConville, M.J., and Billman-Jacobe, H. (2006). Identification of a novel protein with a role in lipoarabinomannan biosynthesis in mycobacteria. J. Biol. Chem. 281, 9011–9017.

43. Marland, Z., Beddoe, T., Zaker-Tabrizi, L., Lucet, I.S., Brammananth, R., Whisstock, J.C., Wilce, M.C.J., Coppel, R.L., Crellin, P.K., and Rossjohn, J. (2006). Hijacking of a substrate-binding protein scaffold for use in mycobacterial cell wall biosynthesis. J. Mol. Biol. 359, 983–997.

44. Yuan, Y., Barrett, D., Zhang, Y., Kahne, D., Sliz, P., and Walker, S. (2007). Crystal structure of a peptidoglycan glycosyltransferase suggests a model for processive glycan chain synthesis. Proc. Natl. Acad. Sci. U. S. A. 104, 5348–5353.

45. Tan, Y.Z., Zhang, L., Rodrigues, J., Zheng, R.B., Giacometti, S.I., Rosário, A.L., Kloss, B., Dandey, V.P., Wei, H., Brunton, R., et al. (2020). Cryo-EM structures and regulation of arabinofuranosyltransferase AftD from mycobacteria. Mol. Cell 78, 683–699.e11.

46. Skovierová, H., Larrouy-Maumus, G., Zhang, J., Kaur, D., Barilone, N., Korduláková, J., Gilleron, M., Guadagnini, S., Belanová, M., Prevost, M.-C., et al. (2009). AftD, a novel essential arabinofuranosyltransferase from mycobacteria. Glycobiology 19, 1235–1247.

47. Kinniment-Williams, B.E., Jurgeleviciute, V., West, D.T., Herman, R., Blaza, J.N., van der Woude, M.W., and Thomas, G.H. (2025). Structural unification of diverse transmembrane acyltransferases reveals a conserved fold for the transmembrane acyl transferase (TmAT) superfamily. J. Biol. Chem. 301, 110546.

48. Yamaryo-Botte, Y., Rainczuk, A.K., Lea-Smith, D.J., Brammananth, R., van der Peet, P.L., Meikle, P., Ralton, J.E., Rupasinghe, T.W.T., Williams, S.J., Coppel, R.L., et al. (2015). Acetylation of trehalose mycolates is required for efficient MmpL-mediated membrane transport in Corynebacterineae. ACS Chem. Biol. 10, 734–746.

49. Belardinelli, J.M., Yazidi, A., Yang, L., Fabre, L., Li, W., Jacques, B., Angala, S.K., Rouiller, I., Zgurskaya, H.I., Sygusch, J., et al. (2016). Structure-function profile of MmpL3, the essential mycolic acid transporter from Mycobacterium tuberculosis. ACS Infect. Dis. 2, 702–713.

50. Belardinelli, J.M., Stevens, C.M., Li, W., Tan, Y.Z., Jones, V., Mancia, F., Zgurskaya, H.I., and Jackson, M. (2019). The MmpL3 interactome reveals a complex crosstalk between cell envelope biosynthesis and cell elongation and division in mycobacteria. Sci. Rep. 9, 10728.

51. Gupta, K.R., Gwin, C.M., Rahlwes, K.C., Biegas, K.J., Wang, C., Park, J.H., Liu, J., Swarts, B.M., Morita, Y.S., and Rego, E.H. (2022). An essential periplasmic protein coordinates lipid trafficking and is required for asymmetric polar growth in mycobacteria. Elife 11, e80395.

52. Xu, R., Ning, Y., Ren, F., Gu, C., Zhu, Z., Pan, X., Pshezhetsky, A.V., Ge, J., and Yu, J. (2024). Structure and mechanism of lysosome transmembrane acetylation by HGSNAT. Nat. Struct. Mol. Biol. 31, 1502–1508.

53. Pearson, C., Tindall, S., Potts, J.R., Thomas, G.H., and van der Woude, M.W. (2022). Diverse functions for acyltransferase-3 proteins in the modification of bacterial cell surfaces: This article is part of the Bacterial Cell Envelopes collection. Microbiology 168. 10.1099/mic.0.001146.

54. Rainczuk, A.K., Klatt, S., Yamaryo-Botté, Y., Brammananth, R., McConville, M.J., Coppel, R.L., and Crellin, P.K. (2020). MtrP, a putative methyltransferase in Corynebacteria, is required for optimal membrane transport of trehalose mycolates. J. Biol. Chem. 295, 6108–6119.

55. Bourgeois, G., Létoquart, J., van Tran, N., and Graille, M. (2017). Trm112, a protein activator of methyltransferases modifying actors of the eukaryotic translational apparatus. Biomolecules 7, 7.

56. Létoquart, J., van Tran, N., Caroline, V., Aleksandrov, A., Lazar, N., van Tilbeurgh, H., Liger, D., and Graille, M. (2015). Insights into molecular plasticity in protein complexes from Trm9-Trm112 tRNA modifying enzyme crystal structure. Nucleic Acids Res. 43, 10989–11002.

57. Wang, C., van Tran, N., Jactel, V., Guérineau, V., and Graille, M. (2020). Structural and functional insights into Archaeoglobus fulgidus m2G10 tRNA methyltransferase Trm11 and its Trm112 activator. Nucleic Acids Res. 48, 11068–11082.

58. Li, W., Shi, Y., Zhang, T., Ye, J., and Ding, J. (2019). Structural insight into human N6amt1-Trm112 complex functioning as a protein methyltransferase. Cell Discov. 5, 51.

59. Liger, D., Mora, L., Lazar, N., Figaro, S., Henri, J., Scrima, N., Buckingham, R.H., van Tilbeurgh, H., Heurgué-Hamard, V., and Graille, M. (2011). Mechanism of activation of methyltransferases involved in translation by the Trm112 “hub” protein. Nucleic Acids Res. 39, 6249–6259.

60. van Tran, N., Muller, L., Ross, R.L., Lestini, R., Létoquart, J., Ulryck, N., Limbach, P.A., de Crécy-Lagard, V., Cianférani, S., and Graille, M. (2018). Evolutionary insights into Trm112-methyltransferase holoenzymes involved in translation between archaea and eukaryotes. Nucleic Acids Res. 46, 8483–8499.

61. Dhiman, R.K., Mahapatra, S., Slayden, R.A., Boyne, M.E., Lenaerts, A., Hinshaw, J.C., Angala, S.K., Chatterjee, D., Biswas, K., Narayanasamy, P., et al. (2009). Menaquinone synthesis is critical for maintaining mycobacterial viability during exponential growth and recovery from non-replicating persistence. Mol. Microbiol. 72, 85–97.

62. McDonough, J.A., McCann, J.R., Tekippe, E.M., Silverman, J.S., Rigel, N.W., and Braunstein, M. (2008). Identification of functional Tat signal sequences in Mycobacterium tuberculosis proteins. J. Bacteriol. 190, 6428–6438.

63. Mohamed, W., Sethi, S., Darji, A., Mraheil, M.A., Hain, T., and Chakraborty, T. (2010). Antibody targeting the ferritin-like protein controls Listeria infection. Infect. Immun. 78, 3306–3314.

64. Krawczyk-Balska, A., and Lipiak, M. (2013). Critical role of a ferritin-like protein in the control of Listeria monocytogenes cell envelope structure and stability under β-lactam pressure. PLoS One 8, e77808.

65. Géraud, N., Rivière, C., Falcou, C., Cioci, G., Froment, C., Gervais, V., Marcoux, J., Gilleron, M., Nigou, J., Fabre, E., et al. (2025). Structural Insights into the Protein Mannosyltransferase from Mycobacterium tuberculosis reveal a WW-Domain-Like Protein Motif in Bacteria. Commun. Biol. 8, 1175.

66. Lommel, M., and Strahl, S. (2009). Protein O-mannosylation: conserved from bacteria to humans. Glycobiology 19, 816–828.

67. Bai, L., Kovach, A., You, Q., Kenny, A., and Li, H. (2019). Structure of the eukaryotic protein O-mannosyltransferase Pmt1-Pmt2 complex. Nat. Struct. Mol. Biol. 26, 704–711.

68. Alexander, J.A.N., and Locher, K.P. (2023). Emerging structural insights into C-type glycosyltransferases. Curr. Opin. Struct. Biol. 79, 102547.

69. Chiapparino, A., Grbavac, A., Jonker, H.R., Hackmann, Y., Mortensen, S., Zatorska, E., Schott, A., Stier, G., Saxena, K., Wild, K., et al. (2020). Functional implications of MIR domains in protein O-mannosylation. Elife 9, e61189.

70. Stallings, C.L., Stephanou, N.C., Chu, L., Hochschild, A., Nickels, B.E., and Glickman, M.S. (2009). CarD is an essential regulator of rRNA transcription required for Mycobacterium tuberculosis persistence. Cell 138, 146–159.

71. Hubin, E.A., Fay, A., Xu, C., Bean, J.M., Saecker, R.M., Glickman, M.S., Darst, S.A., and Campbell, E.A. (2017). Structure and function of the mycobacterial transcription initiation complex with the essential regulator RbpA. Elife 6, e22520.

72. Boyaci, H., Chen, J., Jansen, R., Darst, S.A., and Campbell, E.A. (2019). Structures of an RNA polymerase promoter melting intermediate elucidate DNA unwinding. Nature 565, 382–385.

73. Kumar, A., and Karthikeyan, S. (2018). Crystal structure of the MSMEG_4306 gene product from Mycobacterium smegmatis. Acta Crystallogr. F Struct. Biol. Commun. 74, 166–173.

74. Pereira, L.E., Tsang, J., Mrázek, J., and Hoover, T.R. (2011). The zinc-ribbon domain of Helicobacter pylori HP0958: requirement for RpoN accumulation and possible roles of homologs in other bacteria. Microb. Inform. Exp. 1, 1–10.

75. Barta, M.L., Battaile, K.P., Lovell, S., and Hefty, P.S. (2015). Hypothetical protein CT398 (CdsZ) interacts with σ(54) (RpoN)-holoenzyme and the type III secretion export apparatus in Chlamydia trachomatis: Structure/Function Studies of Chlamydial CT398 (CdsZ). Protein Sci. 24, 1617–1632.

76. Francke, C., Groot Kormelink, T., Hagemeijer, Y., Overmars, L., Sluijter, V., Moezelaar, R., and Siezen, R.J. (2011). Comparative analyses imply that the enigmatic Sigma factor 54 is a central controller of the bacterial exterior. BMC Genomics 12, 385.

77. Delbeau, M., Omollo, E.O., Froom, R., Koh, S., Mooney, R.A., Lilic, M., Brewer, J.J., Rock, J., Darst, S.A., Campbell, E.A., et al. (2023). Structural and functional basis of the universal transcription factor NusG pro-pausing activity in Mycobacterium tuberculosis. Mol. Cell 83, 1474–1488.e8.

78. Kang, J.Y., Mishanina, T.V., Bellecourt, M.J., Mooney, R.A., Darst, S.A., and Landick, R. (2018). RNA polymerase accommodates a pause RNA hairpin by global conformational rearrangements that prolong pausing. Mol. Cell 69, 802–815.e5.

79. Germe, T., Bush, N.G., Baskerville, V., Saman, D., Benesch, J., and Maxwell, A. (2024). Rapid, DNA-induced interface swapping by DNA gyrase. 10.7554/elife.86722.2.

80. Hooper, D.C., and Jacoby, G.A. (2016). Topoisomerase inhibitors: Fluoroquinolone mechanisms of action and resistance. Cold Spring Harb. Perspect. Med. 6, a025320.

81. De Jonge, N., Simic, M., Buts, L., Haesaerts, S., Roelants, K., Garcia-Pino, A., Sterckx, Y., De Greve, H., Lah, J., and Loris, R. (2012). Alternative interactions define gyrase specificity in the CcdB family. Mol. Microbiol. 84, 965–978.

82. VanderSluis, B., Costanzo, M., Billmann, M., Ward, H.N., Myers, C.L., Andrews, B.J., and Boone, C. (2018). Integrating genetic and protein-protein interaction networks maps a functional wiring diagram of a cell. Curr. Opin. Microbiol. 45, 170–179.

83. Reid, A.J., Ranea, J.A., and Orengo, C.A. (2010). Comparative evolutionary analysis of protein complexes in E. coli and yeast. BMC Genomics 11, 79.

84. Janes, J. pooled-ppi-yeast: Work-in-progress map of yeast protein-protein interactions using pooled co-folding. https://github.com/jurgjn/pooled-ppi-yeast (Github).

85. Peters, J.M., Koo, B.-M., Patino, R., Heussler, G.E., Hearne, C.C., Qu, J., Inclan, Y.F., Hawkins, J.S., Lu, C.H.S., Silvis, M.R., et al. (2019). Enabling genetic analysis of diverse bacteria with Mobile-CRISPRi. Nat. Microbiol. 4, 244–250.

86. Koo, B.-M., Todor, H., Sun, J., van Gestel, J., Hawkins, J.S., Hearne, C.C., Banta, A.B., Huang, K.C., Peters, J.M., and Gross, C.A. (2025). Comprehensive genetic interaction analysis of the Bacillus subtilis envelope using double-CRISPRi. Cell Syst. 16, 101406.

87. Dénéréaz, J., Eray, E., Jana, B., de Bakker, V., Todor, H., van Opijnen, T., Liu, X., and Veening, J.-W. (2025). Dual CRISPRi-seq for genome-wide genetic interaction studies identifies key genes involved in the pneumococcal cell cycle. Cell Syst. 16, 101408.

88. Kapopoulou, A., Lew, J.M., and Cole, S.T. (2011). The MycoBrowser portal: a comprehensive and manually annotated resource for mycobacterial genomes. Tuberculosis (Edinb.) 91, 8–13.

89. Kim, L.M., Todor, H., and Gross, C.A. (2024). Correction of a widespread bias in pooled chemical genomics screens improves their interpretability. Mol. Syst. Biol. 20, 1173–1186.

90. Pelicic, V., Jackson, M., Reyrat, J.M., Jacobs, W.R., Jr, Gicquel, B., and Guilhot, C. (1997). Efficient allelic exchange and transposon mutagenesis in Mycobacterium tuberculosis. Proc. Natl. Acad. Sci. U. S. A. 94, 10955–10960.

91. Jackson, M., Crick, D.C., and Brennan, P.J. (2000). Phosphatidylinositol is an essential phospholipid of mycobacteria. J. Biol. Chem. 275, 30092–30099.

92. Méderlé, I., Bourguin, I., Ensergueix, D., Badell, E., Moniz-Peireira, J., Gicquel, B., and Winter, N. (2002). Plasmidic versus insertional cloning of heterologous genes in Mycobacterium bovis BCG: impact on in vivo antigen persistence and immune responses. Infect. Immun. 70, 303–314.

93. Murphy, K.C., Nelson, S.J., Nambi, S., Papavinasasundaram, K., Baer, C.E., and Sassetti, C.M. (2018). ORBIT: A new paradigm for genetic engineering of Mycobacterial chromosomes. MBio 9. 10.1128/mbio.01467-18.

94. Frejno, M., Berger, M.T., Tüshaus, J., Hogrebe, A., Seefried, F., Graber, M., Samaras, P., Ben Fredj, S., Sukumar, V., Eljagh, L., et al. (2025). Unifying the analysis of bottom-up proteomics data with CHIMERYS. Nat. Methods 22, 1017–1027.

